# Repeat associated non-AUG translation as a common mechanism for the polyGln ataxias

**DOI:** 10.1101/2025.10.14.682372

**Authors:** Monica Banez Coronel, Tao Zu, Madeline Denton, Shu Guo, Ramadan Ajredini, Deborah Morrison, Alexis B. Tays, Olga Pletnikova, Anthony T. Yachnis, Juan C. Troncoso, Henry L. Paulson, Hayley S. McLoughlin, Tetsuo Ashizawa, S.H. Subramony, Laura P.W. Ranum

## Abstract

Determining if repeat associated non-AUG (RAN) proteins contribute to the CAG polyGln-encoding spinocerebellar ataxias (CAG-SCAs) is critical for understanding mechanisms and developing therapies for these diseases. Immunohistochemistry using antibodies against polySer and polyLeu repeats and locus specific C-terminal regions show sense polySer (AGC frame) and antisense polyLeu (CUG frame) RAN proteins accumulate in affected grey and white-matter brain regions, throughout the cerebellum and pons, in SCA1, SCA2, SCA3, SCA6, and SCA7 autopsy brains. Cerebellar white matter regions with prominent polySer and polyLeu but minimal polyGln aggregates show demyelination, white matter loss, and activated microglia. In SCA3 mice, RAN proteins accumulate in an age-dependent manner. In neural cells, polySer and polyLeu RAN proteins are toxic and cause autophagic dysfunction. In cells, the FDA-approved drug metformin decreases RAN protein levels and reduces toxicity. Taken together, these data identify sense and antisense RAN proteins as a common molecular mechanism shared by the CAG-SCAs.

## INTRODUCTION

The polyglutamine (polyGln) encoding spinocerebellar ataxias (SCAs) are a group of dominantly inherited repeat expansion diseases that cause neurodegeneration in the cerebellum and brainstem. Among these, SCA1, SCA2, SCA3, SCA6, and SCA7 are caused by CAG•CTG repeat expansion mutations in which the CAG expansion mutations encode long polyGln tracts within their respective and diverse proteins^1,2^. Although the proteins encoded by each of these genes have different functions, these progressive neurodegenerative disorders primarily affect the cerebellum and brainstem, which in turn leads to balance, speech, and eye movement abnormities. While substantial insights have been made in understanding the clinical, pathological, and molecular features of the polyGln SCAs, our understanding of these diseases is incomplete and no effective treatment strategies are available for any of these disorders.

Regardless of the position of the mutation within its respective gene, a growing number of microsatellite expansion mutations produce both sense and antisense expansion transcripts^3–10^. Additionally, microsatellite expansion mutations can be translated in all three frames in the absence of an AUG- or AUG-like initiation codons^11^. This repeat associated non-AUG (RAN) translation, was initially reported in SCA8 and myotonic dystrophy type 1 (DM1)^11^ and subsequently in other non-coding expansion disorders (e.g. C9orf72 Amyotrophic lateral sclerosis (ALS) and fragile-X associated tremor and ataxia syndrome [FXTAS]^11–14^). In 2015, sense and antisense RAN proteins were also reported in the polyGln disorder Huntington’s disease (HD)^15^. RAN protein aggregates have been previously reported in patient tissues in 11 different diseases^16,17^.

Evidence that RAN proteins are toxic has been shown in multiple studies^8,11,15,18–26^. For example, in *C9orf72* ALS/FTD mice, treatment with α-GA RAN protein antibodies^18^ or decreasing RAN protein levels by inhibition of the protein kinase R (PKR) pathway^21^ showed improved behavior, decreased neuroinflammation and increased motor neuron survival. Immunohistochemistry (IHC) studies performed on autopsy brains from myotonic dystrophy type 2 (DM2) patients show RAN protein aggregates preferentially accumulate in necrotic brain regions^22^. Similarly, in SCA8 autopsy tissue, polySer RAN protein aggregates are found in white matter regions showing demyelination and axonal breakage^27^. In HD, sense and antisense RAN proteins are abundantly found in regions with activated caspase-3 staining, activated microglia, and astrogliosis^15^.

A major obstacle in determining if RAN proteins contribute to the CAG•SCAs has been the lack of tools to detect shared homopolymeric repeats across diseases. The original report showing that RAN proteins accumulate in HD used locus-specific C-terminal (Ct) antibodies^15^. While useful and important for locus specific confirmation, disease specific C-terminal antibodies such as those used to study RAN proteins in HD cannot be used to detect RAN protein accumulation in other CAG•CTG expansion disorders.

Using a combination of newly developed polySer and polyLeu repeat motif and locus-specific C-terminal antibodies, we show polySer (sense) and polyLeu (antisense) RAN proteins aggregates are a shared pathologic feature found in affected SCA1, SCA2, SCA3, SCA6, and SCA7 autopsy brains. RAN protein aggregates prominently accumulate in vulnerable brain regions including damaged white matter regions of the cerebellum. RAN proteins are found in SCA1 and SCA3 mice. Time course studies using SCA3 mice show RAN protein aggregates increase with age. Alternative codon experiments show locus specific SCA1, SCA2, SCA3, SCA6 and SCA7 polySer and polyLeu RAN proteins are toxic to neural cells. Additionally, decreasing RAN protein levels with metformin reduces cell toxicity.

## RESULTS

### Validation of novel α-polySer and α-polyLeu repeat antibodies

To explore the contribution of RAN translation across the CAG-SCAs we developed custom polyclonal antibodies directed against polySer (sense AGC frame) and polyLeu (antisense CUG frame) repeat motifs (**Figure 1A**). Antibody specificity was tested in HEK293T cells expressing minigenes containing an N-terminal ATG-initiated flag epitope tag followed by expanded polySer or polyLeu repeat motifs (**Figure 1B-C**). Immunofluorescence (IF) experiments in transfected HEK293T cells show colocalization between α-polySer or α-polyLeu and α-flag-epitope signals (**Figure 1D-E**). The ability of these novel antibodies to detect polySer and polyLeu RAN proteins in human postmortem tissue was tested using RAN positive SCA8 and HD autopsy brains. IHC experiments using our novel α-polySer antibody showed nuclear polySer staining similar to that found with our previously validated α-SCA8-Ser-Ct and α-HD-Ser-Ct antibodies in SCA8 and HD autopsy brains, respectively **(Figures 1F, S1A-B**). Similarly, IHC staining using our novel α-polyLeu antibody showed distinct perinuclear polyLeu aggregates in HD autopsy brains consistent with staining using our previously validated HD-Leu-Ct antibody (**Figures 1G, S1C**). No similar staining was detected with either antibody in controls or in HD tissue stained with α-polySer or α-polyLeu pre-immune sera (**Figures 1, S1**). These results demonstrate that novel α-polySer and α-polyLeu repeat antibodies are specific and sensitive tools to explore RAN protein accumulation across CAG•CTG expansion disorders.

**Figure 1.**
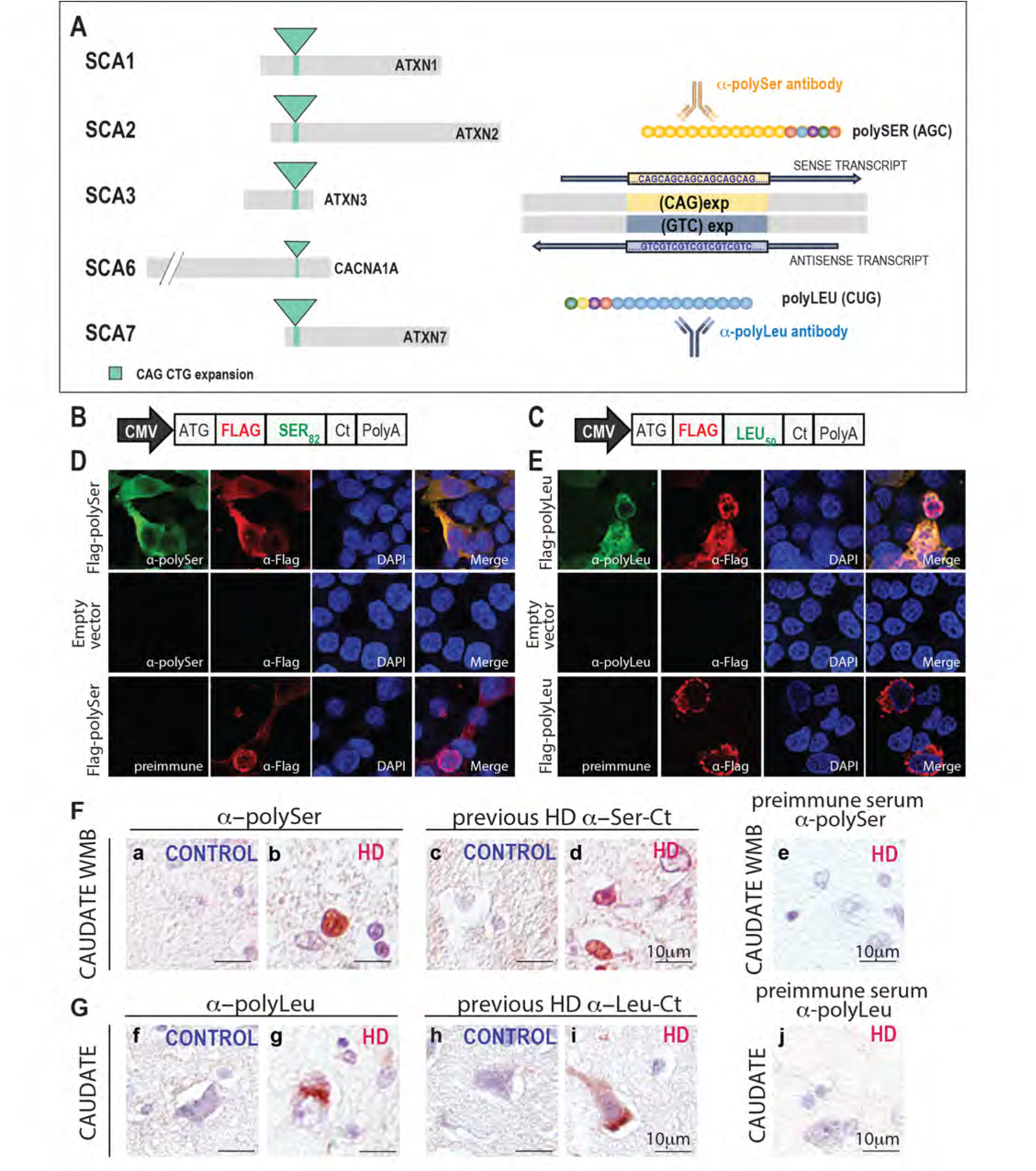
Validation of novel α-polySer and α-polyLeu antibodies. (A) Schematic diagram depicting polyGln encoding genes and their corresponding repeat expansion mutations in the polyGln-SCAs (left) and antibodies designed to test if sense-encoded polySer and antisense-encoded polyLeu RAN proteins are expressed across these polyGln encoding CAG expansion mutations. (B-C) Constructs used to express Flag-polySer and Flag-polyLeu expansion proteins. (D-E) Representative IF images showing co-localization of signal from newly developed **α**-polySer or **α**-polyLeu repeat antibodies (green) and α-Flag (red) in transfected HEK293T cells (upper panels). No polySer or polyLeu signal was detected in cells transfected with empty vector (center) or pre-immune serum (lower panels). (F) Our newly developed α-polySer antibody detects nuclear staining (b) in representative images from white matter striatal bundles similar to staining using a previously validated HD-Ser-Ct antibody^15^ (d) in HD but not unaffected controls (a,c) or preimmune serum (e) (G) Similarly, IHC using newly developed α-polyLeu and previously validated HD-Leu-Ct antibodies show similar representative staining in HD (g,i) but not control caudate (f,h) or HD tissue stained with pre-immune sera (j). Red, positive staining; purple, counterstain; WMB, white matter bundles.

### RAN aggregates are a common pathological feature across the CAG•CTG polyGln SCAs

#### Sense polySer RAN proteins

Human autopsy brains from patients diagnosed with spinocerebellar ataxia were collected as part of the University of Florida / National Ataxia Brain Bank and genetically tested to confirm genetic diagnosis (**Table S1**). IHC using our validated α-polySer antibody detected polySer aggregates in genetically confirmed SCA1 (n=5), SCA2 (n=5), SCA3 (n=6), SCA6 (n=4) and SCA7 (n=5) human autopsy brains (**Figure 2**, **Table 1**). Robust α-polySer staining was seen throughout the cerebellum with the most prominent signal detected in white matter (WM) regions of the cerebellum, the deep cerebellar dentate nuclei (DCN), and Bergmann glia surrounding the Purkinje cells (**Figures 2A, S2A, Table 1**). No similar staining was detected in SCA5 (n=2), which is not caused by a repeat expansion, or unaffected control (n=5) brains. In the granule cell layer (GCL), RAN protein staining is found in each of the polyGln SCAs but is variable between cases. Some cases show intense soluble and aggregate staining (e.g. SCA2 and SCA6, **Figure S2A**) in the GCL while other cases show more predominant GCL aggregates (e.g. SCA1 **Figure S2A**). In the DCN, polySer aggregates are detected in dentate neurons, glia, and throughout the neuropil (**Figures 2A, S2A**). Similarly, the pons, which can be significantly affected in the polyGln SCAs, shows frequent and robust RAN polySer staining in neurons and glia in SCA1, SCA2, SCA3, SCA6, and SCA7, but not controls (**Figures 2B, S2B**). In the less affected frontal cortex, polySer staining is rare. When staining is detected, it usually presents as a diffuse faint signal with occasional microaggregates. No similar signal was detected in unaffected or SCA5 controls (**Figures 2C, S2C**). Although the abundance of polySer staining varies among individuals, distinct disease-specific distribution patterns are observed across the polyGln SCAs (**Table 1**). For example, prominent polySer accumulation in Bergmann glia is frequently observed in SCA2, SCA6, and SCA7 cases, but was less frequently seen in the SCA3 cases examined (**Table 1**). In the deep white matter regions (DWM) containing the dentate nuclei, neuronal polySer aggregates were more frequently found in SCA2, SCA6 and SCA7 compared to SCA1 and SCA3.

**Figure 2.**
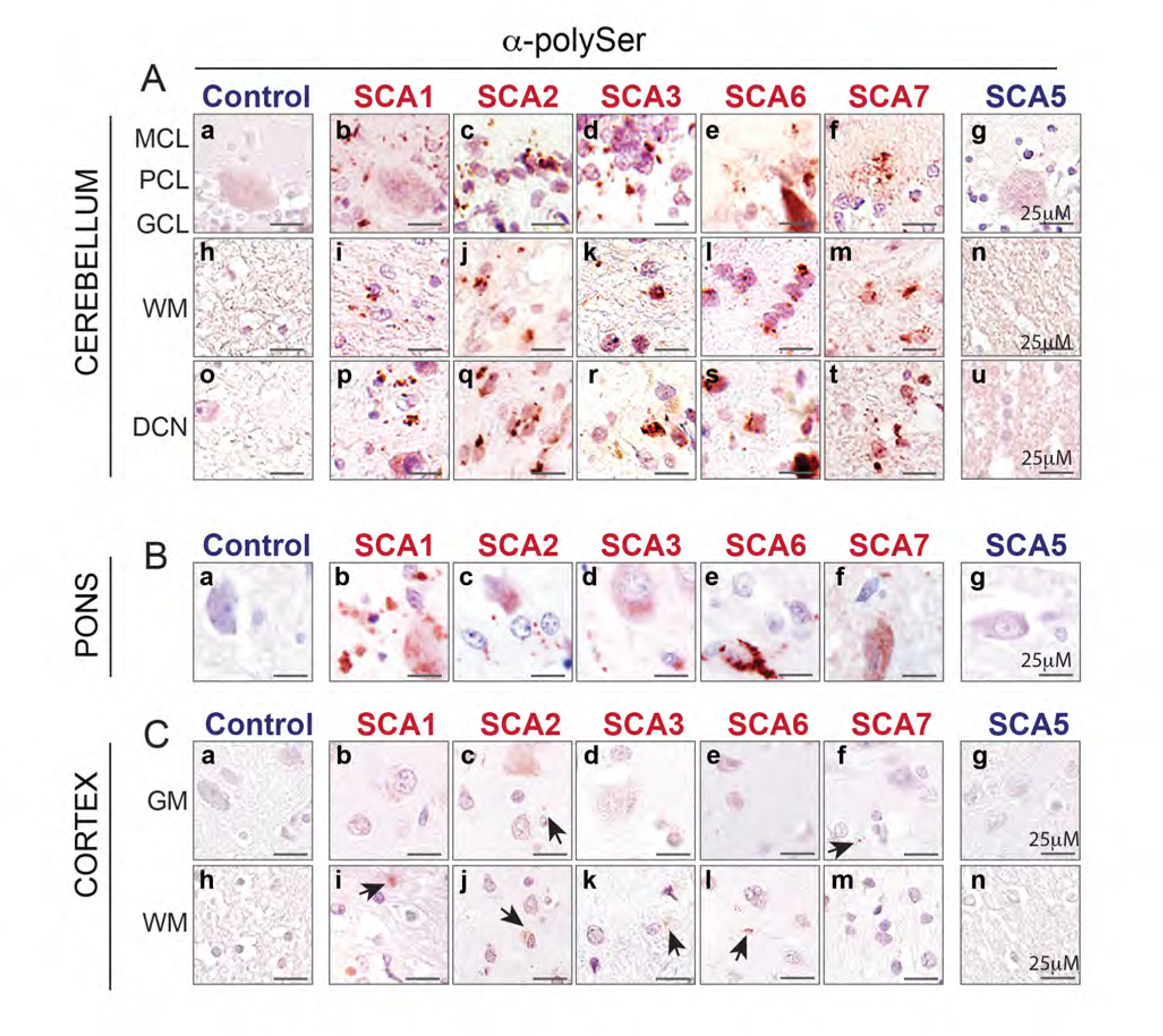
Sense RAN polySer aggregates in SCA1, SCA2, SCA3, SCA6, and SCA7 cerebellum and pons. (A) Representative IHC images show prominent α-polySer positive aggregates in cerebellum from SCA1, SCA2, SCA3, SCA6, and SCA7 autopsy brains. Positive staining is found in Bergmann glia surrounding the Purkinje cells (a-g), white matter (h-n), and deep white matter regions around the dentate nuclei (o-u) in the polyGln-SCAs but not in tissue from unaffected or SCA5 disease controls. (B) In pons, frequent polySer aggregates accumulate in neurons and glia in the CAG-SCAs but not controls (a-g). (C) In contrast, in the frontal cortex α-polySer staining in the CAG-SCAs is rare in both grey (b-f) and white matter (i-m) regions. GCL, granular cell layer; PCL, Purkinje cell layer; MCL, molecular cell layer; DCN, deep cerebellar nuclei; WM, white matter; GM, grey matter. Red, positive staining; purple, nuclear counterstain. SCA1, n=5; SCA2, n=5; SCA3, n=6; SCA6, n=4; SCA7, n=5; unaffected controls (n=5); SCA5 disease controls, n=2.

**Table 1.**
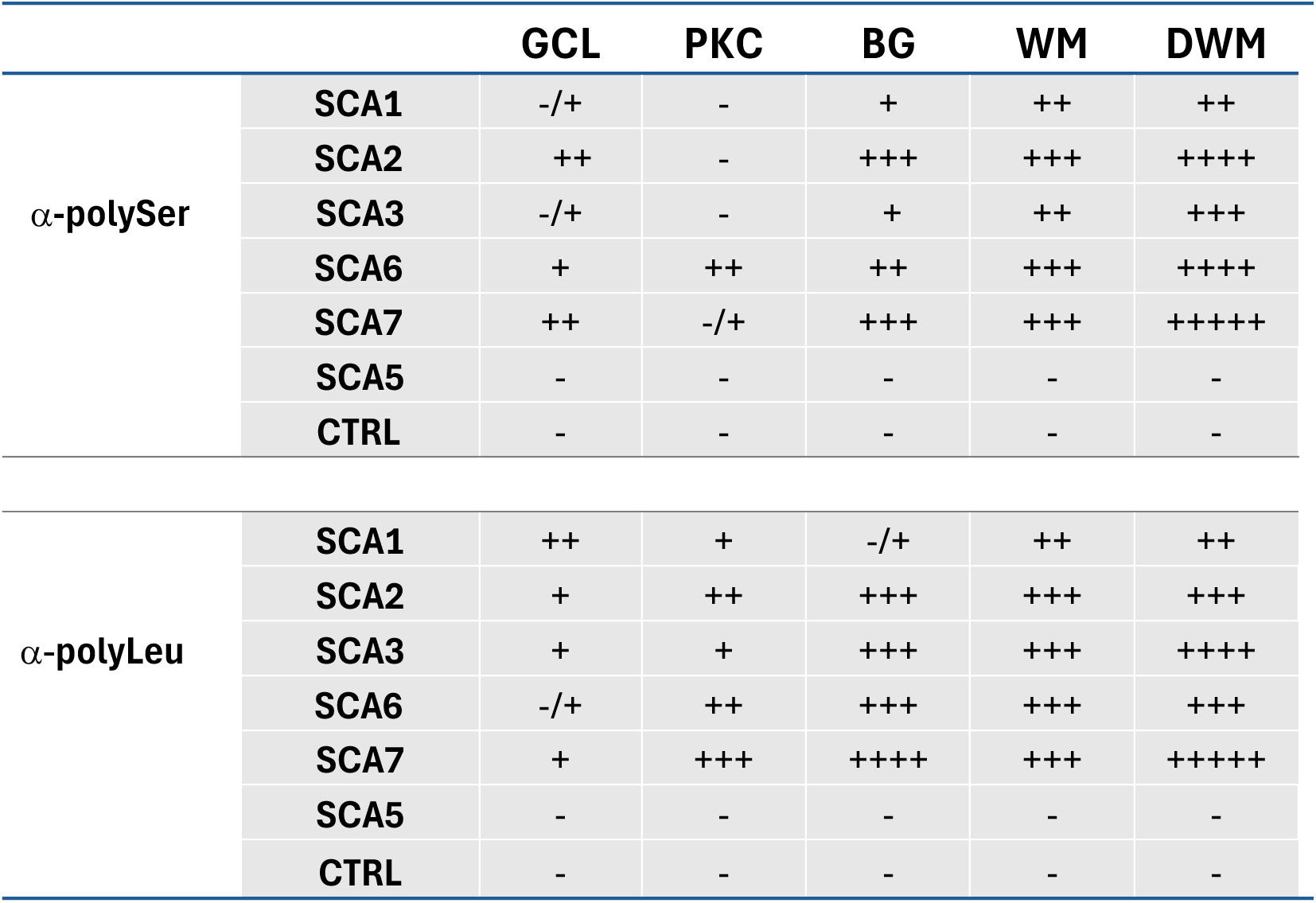
Summary of α-polySer and α-polyLeu RAN protein staining in SCA1, SCA2, SCA3, SCA6 and SCA7 cerebellar regions. RAN staining: -, negative; -/+, rare low-intensity staining; +, low-intensity staining; ++, moderate intensity; +++, frequent and intense; ++++, very frequent positive cells with high-intensity staining, +++++ Robust and intense staining in most cells (>85% positive cells). GCL=granule cell layer, PKC=Purkinje cells, BG=Bergman glia, WM=white matter, DWM=deep white matter

#### Anti-sense PolyLeu RAN proteins

Because bidirectional transcription and sense and antisense RAN proteins have been reported in a number of expansion disorders, we tested if antisense polyLeu RAN proteins accumulate across the CAG-SCAs. IHC using our novel α-polyLeu antibody detected frequent polyLeu aggregates in the Bergmann glia, cerebellar cortical white matter, and deep white matter regions of the DCN (**Figures 3A, S3A, Table 1**). In contrast to polySer aggregates, polyLeu aggregates were more intense and more frequently found in Purkinje cells and dentate neurons in the DCN region (**Figures 3A, S3A, Table 1).** In the pons, α-polyLeu staining showed frequent, robust aggregates in the neuropil and glia. In contrast, polyLeu staining was more diffuse in pontine neurons (**Figures 3B, S3B**). In the less affected frontal cortex, polyLeu staining was rare (**Figures 3C, S3C**).

**Figure 3.**
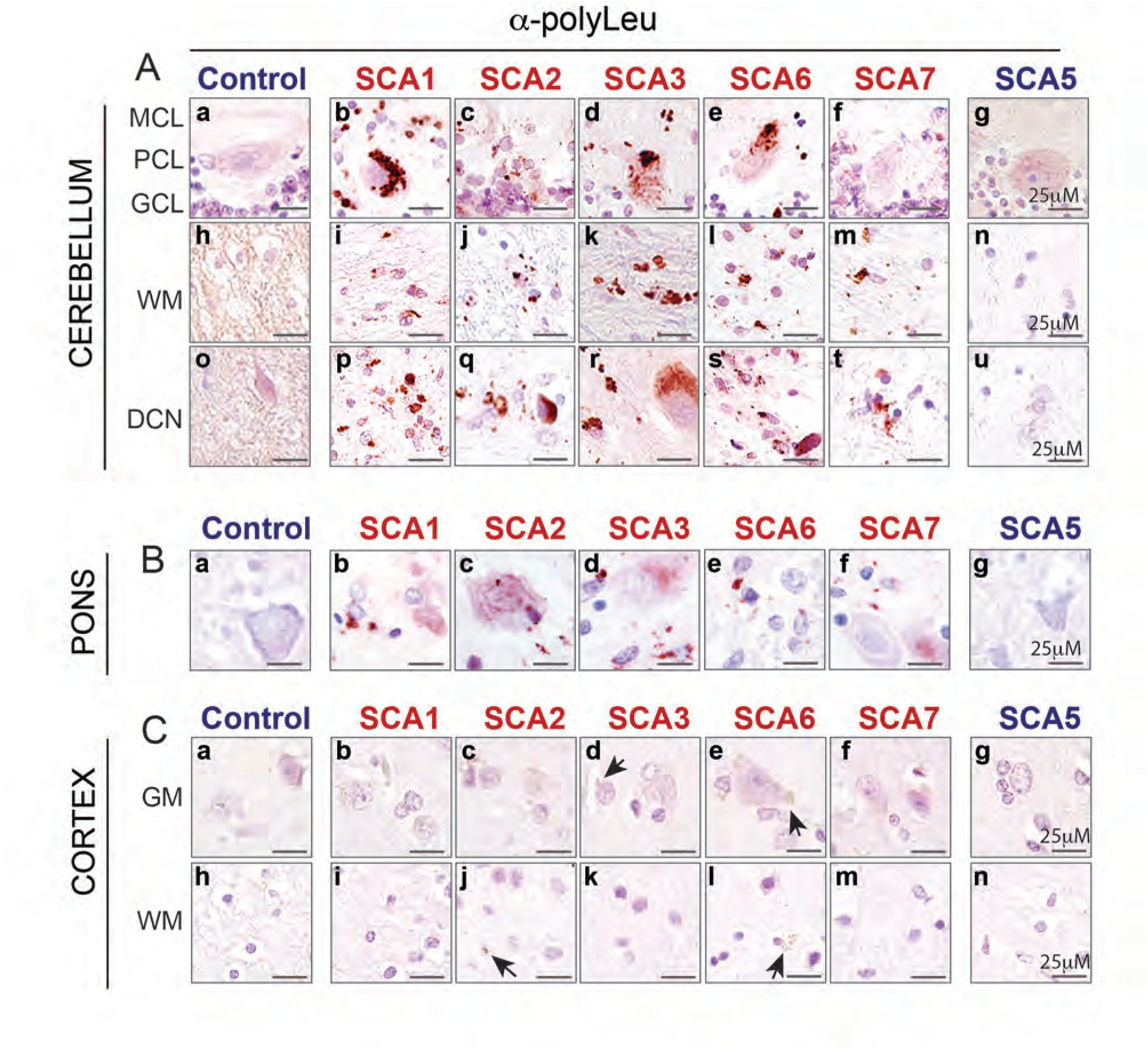
Antisense RAN polyLeu aggregates in SCA1, SCA2, SCA3, SCA6, and SCA7 cerebellum and pons. (A) α-polyLeu immunostaining shows frequent and robust aggregate staining throughout the cerebellum in SCA1, SCA2, SCA3, SCA6, and SCA7 but not unaffected or SCA5 disease controls. PolyLeu positive cells include Purkinje cells, neurons of the DCN and glial cells. (B) Similar α-polyLeu staining is also found in pons across the CAG-SCAs, but not controls. (C) In contrast, only very rare and much lighter α-polyLeu staining was observed in the grey (a-g) and white matter regions (h-n) of the frontal cortex in the CAG-SCAs but not controls. GCL, granular cell layer; PCL, Purkinje cell layer; MCL, molecular cell layer; DCN, deep cerebellar nuclei. WM, white matter; GM, grey matter Red, positive staining; purple, nuclear counterstain. SCA1, n=5; SCA2, n=5; SCA3, n=6; SCA6, n=4; SCA7, n=5; unaffected controls (n=5); SCA5 disease controls, n=2.

In summary, these data identify sense polySer and antisense polyLeu RAN proteins as novel and frequent histopathological signatures found in affected brain regions across the polyGln CAG-SCAs.

### Locus-specific C-terminal antibodies confirm polySer and polyLeu proteins are expressed from SCA1, SCA2, SCA3, SCA6, or SCA7 expansion mutations

To test if staining using the α-polySer and α-polyLeu antibodies detects RAN proteins expressed from expanded repeats at the SCA1, SCA2, SCA3, SCA6, or SCA7 loci, we generated locus-specific antibodies against the unique Ct regions of each of these proteins. IHC of serial sections confirms that α-polySer and α-polyLeu positive samples also showed positive staining for their corresponding locus-specific Ct-antibody (**Figure 4**). For example, cerebellar tissue from SCA1 cases positive for α-polySer, also showed positive staining with the SCA1 locus-specific C-terminal antibody but not the other locus specific C-terminal antibodies (i.e. α-SCA2-Ser-Ct, α-SCA3-Ser-Ct, α-SCA6-Ser-Ct, or α-SCA7-Ser-Ct). Similarly, cerebellar tissue from SCA7 patients positive for α-polyLeu were also positive for the α-SCA7-Leu-Ct but negative for α-SCA1-Leu-Ct, α-SCA2-Leu-Ct, α-SCA3-Leu-Ct and α-SCA6-Leu-Ct) (**Table S2**). The only notable exceptions were found in individual SCA1, SCA3, SCA6 and SCA7 cases that showed rare but positive α-SCA2-Leu-Ct aggregate staining, not typical of SCA2 cases (**Table S2**). All SCA cases were genetically confirmed as SCA1, SCA2, SCA3, SCA6 or SCA7 with no cases showing more than one mutation, or premutation alleles. While premutation alleles at the SCA2 locus do not explain the rare staining seen in a subset of non-SCA2 patients (**Table S2**), it is possible that the presence of a close cognate AAG codon in the polyLeu reading frame may allow occasional readthrough and expression of SCA2 polyLeu proteins from these normal length alleles.

**Figure 4.**
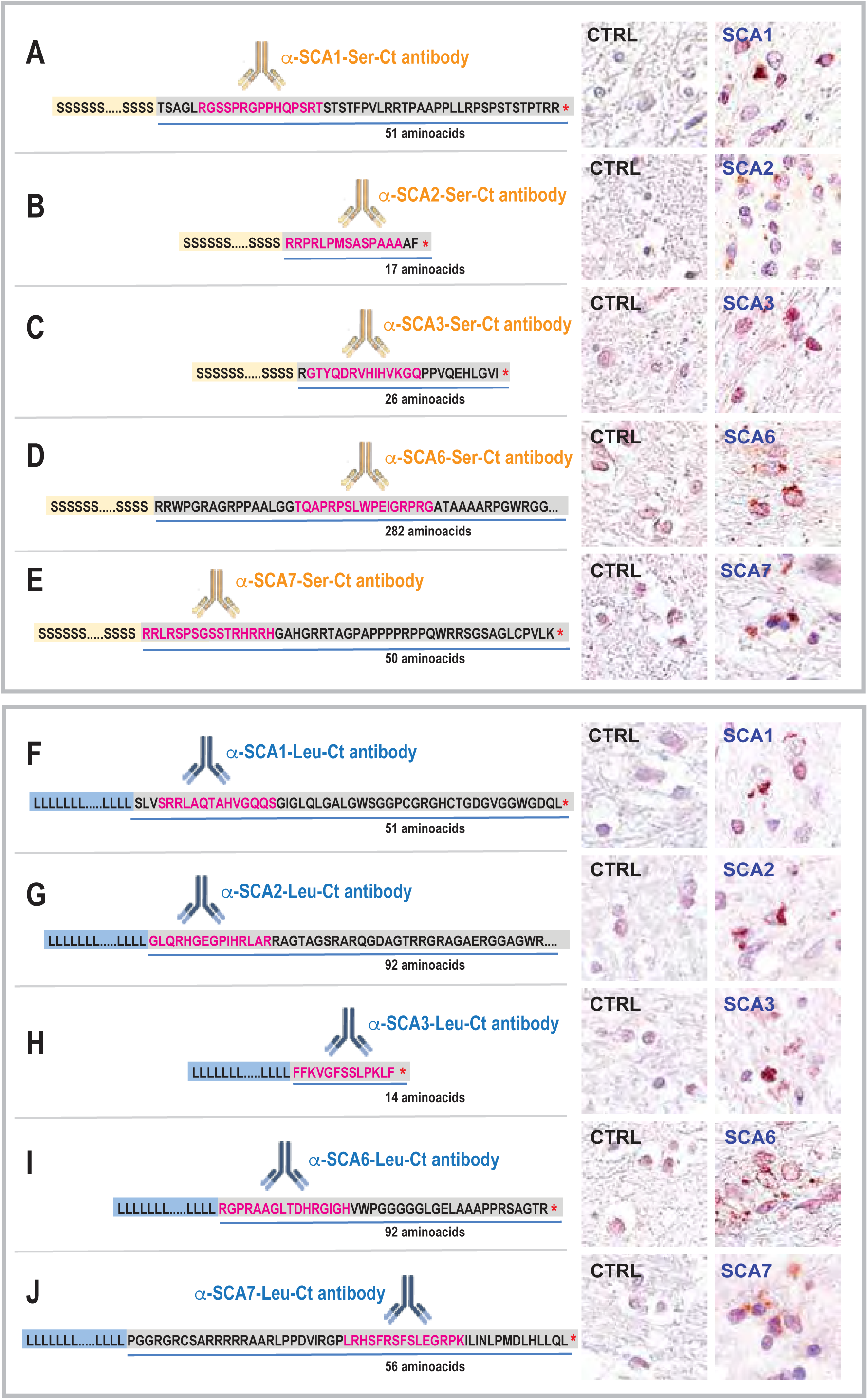
PolySer and polyLeu RAN proteins detected using independent locus-specific SCA1, SCA2, SCA3, SCA6, and SCA7 C-terminal antibodies. (Left column) Schematic diagram showing putative CAG-SCA RAN polySer (A-E) and polyLeu (F-J) proteins with unique C-terminal (Ct) regions. Peptide sequences used to generate locus-specific Ct antibodies are shown in magenta (Right) IHC staining with locus-specific antibodies demonstrate that polySer (A-E) and polyLeu (F-J) proteins expressed from the SCA1, SCA2, SCA3, SCA6, and SCA7 mutations accumulate as aggregates in affected cerebellum.

While SCA1, SCA2, SCA3, SCA6, and SCA7 all express polySer and polyLeu containing expansion proteins that form aggregates in vulnerable brain regions, each of these proteins contains unique C-terminal regions that substantially vary in both sequence and length (**Figure 4**).

### Abundant PolySer and polyLeu aggregates in white matter regions with demyelination and neuroinflammation

White matter abnormalities have been reported in SCA1, SCA2, SCA3, SCA6, and SCA7^28–30^ and linked to disease severity in SCA2, SCA3, and SCA7^31–35^. Interestingly, white matter regions show only very rare and infrequent polyGln aggregates. To test if RAN protein aggregates could explain the histopathological changes found in brain regions with white matter abnormalities, we performed a series of comparative histological studies on cerebellum and frontal cortex using α-polySer and α-polyLeu, 1C2 (polyGln), α-Iba1 (microglia) and luxol fast blue (LFB, demyelination) staining. IHC of serial sections shows prominent α-polySer and α-polyLeu staining in cerebellar white matter regions with demyelination and white matter loss in SCA1, SCA2, SCA3, SCA6, and SCA7 (**Figures 5A-B, S4A, Table 2**). RAN-positive regions also have increased numbers of astrocytes (determined by nuclear morphology) and microglia, which often show a round amoeboid morphology indicative of neuroinflammation and active phagocytosis (**Figures 5A-B, S4A, Table 2**)^36–39^. Consistent with previous studies, polyGln aggregates in these white matter regions are extremely rare. While prominent RAN protein pathology is found in damaged cerebellar white matter, RAN protein aggregates are rare in the less affected white matter regions of the frontal cortex, which shows minimal demyelination and microglial proliferation/activation (**Figures 5A**, **5C, S4B, Table 2**).

**Figure 5.**
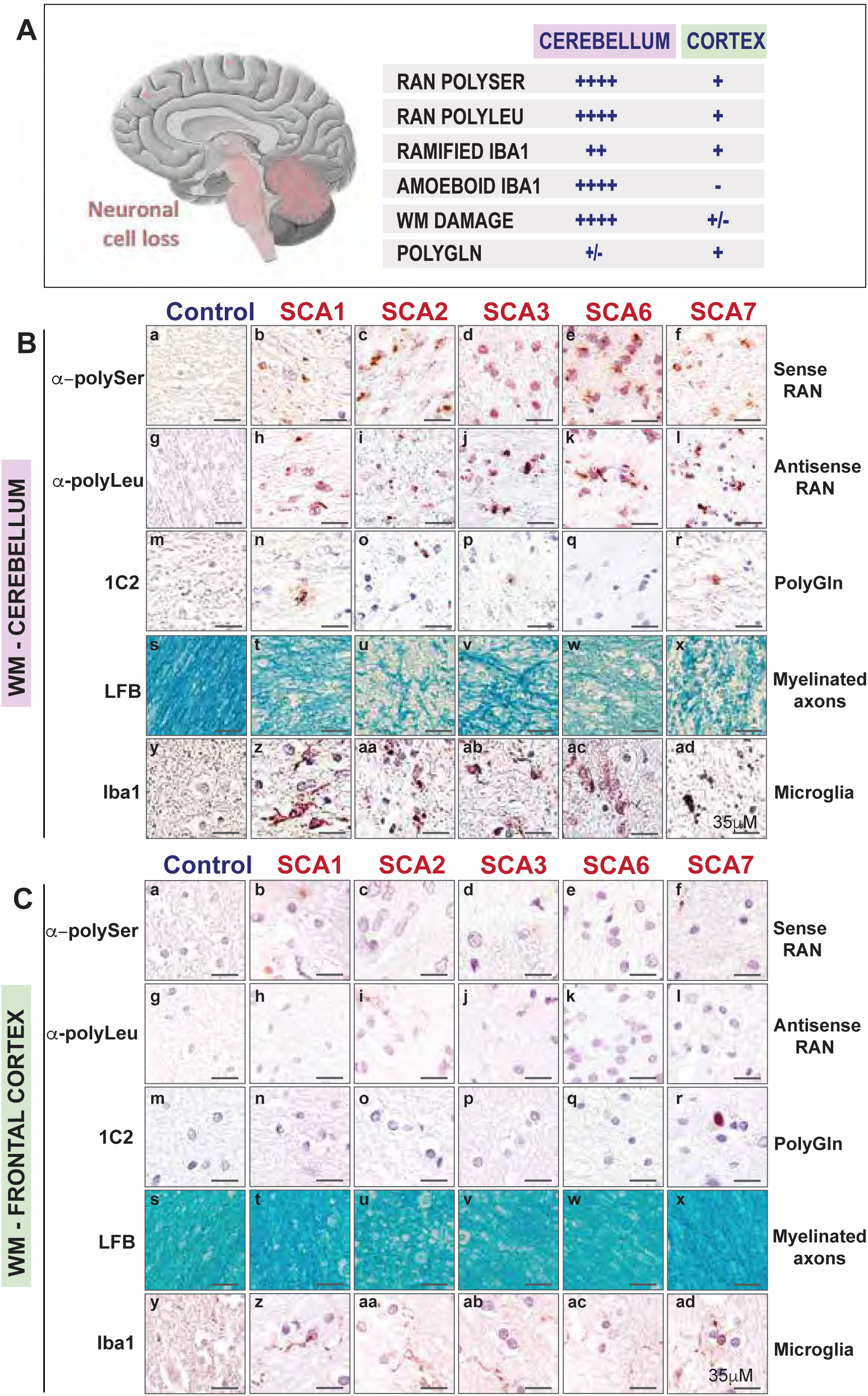
PolySer and polyLeu RAN proteins accumulate in white matter cerebellar regions showing demyelination and neuroinflammatory changes. (A) Schematic diagram showing brain regions affected in the CAG-SCAs and table summarizing the IHC findings. (B) IHC using serial sections of SCA1, SCA2, SCA3, SCA6 and SCA7 cerebellum show frequent RAN polySer (a-f) and RAN polyLeu (g-l) aggregates, but only rare polyGln aggregates (m-r), in white matter regions with prominent demyelination (s-x, decreased LFB signal) and increased microglia (y-ad) compared to controls. (C) White matter regions in the frontal cortex have minimal polyGln and polySer and polyLeu RAN proteins with no differences in LFB or Iba1 staining compared to controls. LFB, luxol fast blue. WM, white matter. Red, positive staining; purple, nuclear counterstain. Control, n=5, SCA1, n=5; SCA2, n=5; SCA3, n=6; SCA6, n=4; SCA7, n=5.

**Table 2.**
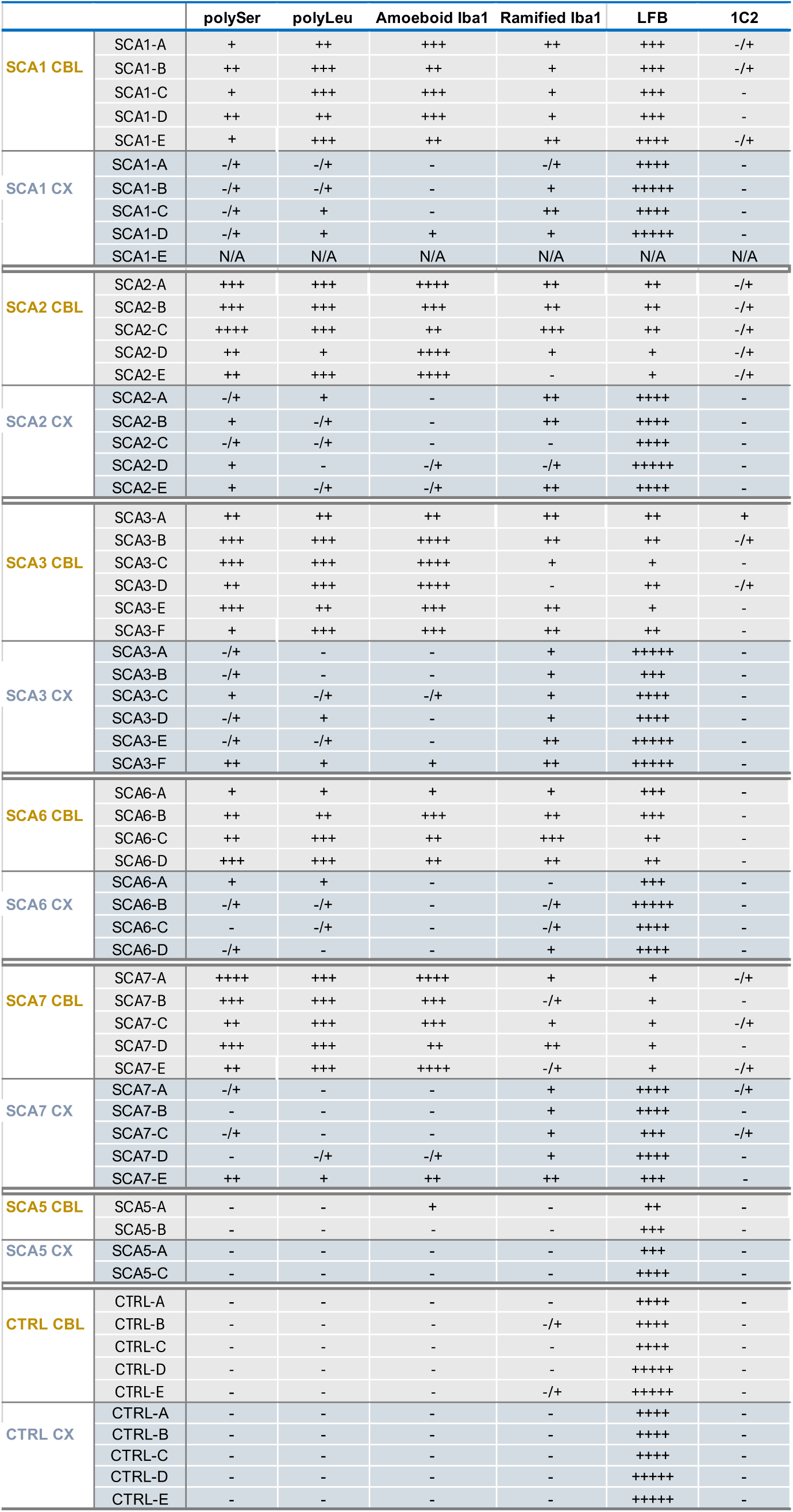
Summary of RAN protein histopathology in cerebellum and frontal cortex in individual SCA cases and controls. RAN staining: -, negative; -/+, rare low intensity staining; +, low-intensity staining; ++, moderate intensity; +++, frequent and intense; ++++, very frequent positive cells with high intensity staining, +++++ Robust and intense staining in most cells (>85% positive cells); N/A, not available.

Taken together, these data suggest that RAN proteins are an important driver of cerebellar white matter pathology across the CAG•CTG polyGln ataxias.

### Sense and antisense RAN proteins accumulate in ataxia mouse models

Mouse models have been used for many years to understand the effects of mutant polyGln proteins in the polyGln-encoding SCAs. Using SCA1 and SCA3 as examples we tested the hypothesis that these models also express RAN proteins. We chose SCA1 knock-in (SCA1^154Q/WT^)^40,41^ and SCA3 YAC (SCA3^Q84/WT^)^42^ models because the repeat expansion mutations are located in the context of their corresponding full-length mouse or human genes and endogenous promoter regions. In SCA1 KI mice, RAN polySer and polyLeu aggregates accumulate as nuclear and perinuclear aggregates in the cortex of 22-week old SCA1^154Q/WT^ knock-in mice (**Figure S5A**). Rare aggregates were also detected in cerebellar deep white matter regions and around the dentate nuclei (not shown). In SCA3 YAC mice, sense polySer and antisense polyLeu RAN protein aggregates were found primarily in the granule cell layer close to the Purkinje cell layer, and in the cerebellar white matter and pons (**Figure S5B-C**). Previous longitudinal studies in SCA3 YAC have shown worsening behavioral and neurodegenerative phenotypes with age, which have been proposed to result from increased polyGln aggregates. Here we show sense and antisense RAN proteins also increase with mouse age (**Figure 6)** in cerebellum and pons, which were both previously reported as vulnerable brain regions in these mice^42^.

**Figure 6.**
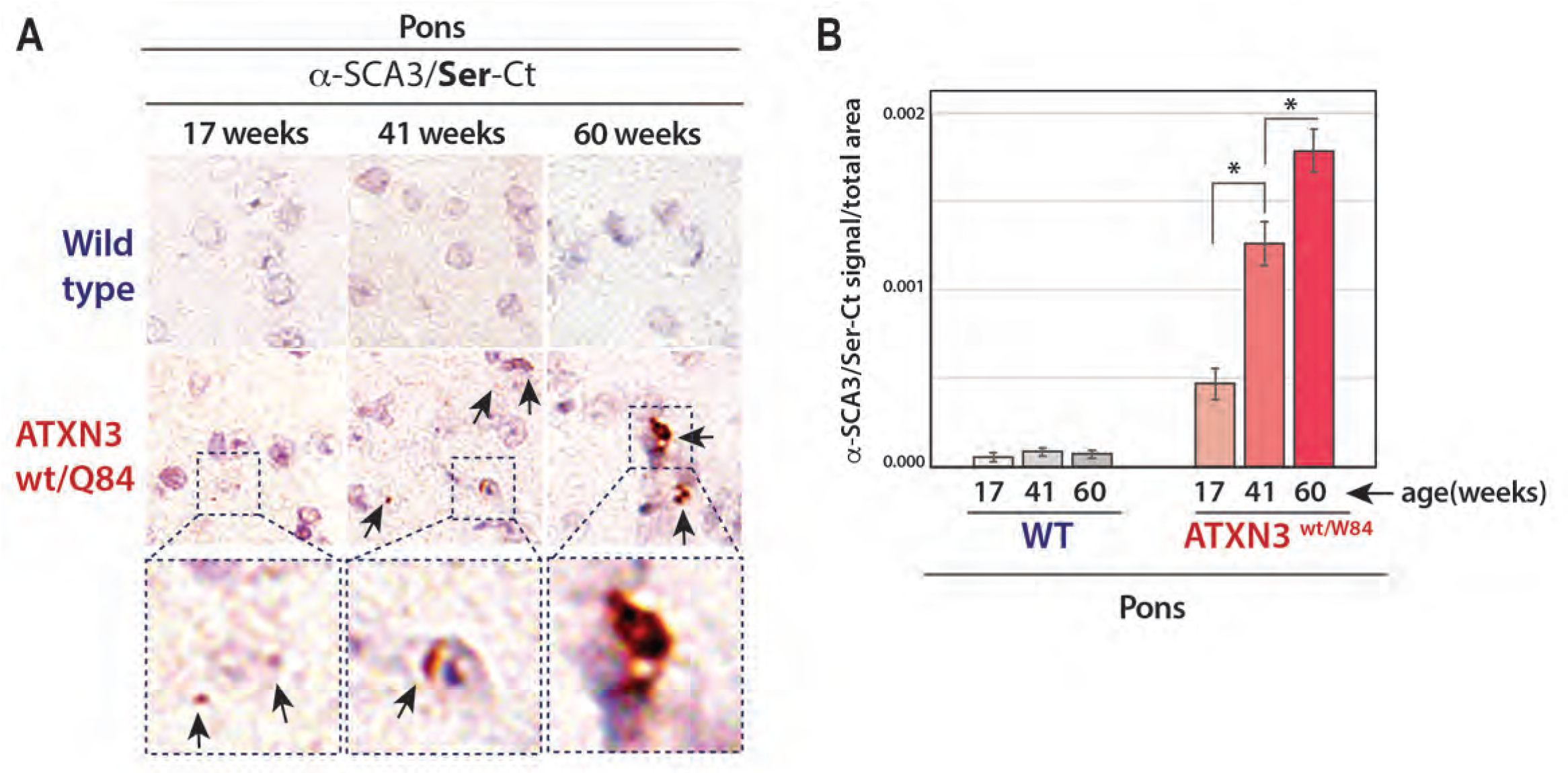
RAN protein aggregates increase with age in SCA3 YAC mice. (A) IHC images show pontine polySer RAN aggregates increase in number and intensity with age in ATNX3^Q84/WT^ mice. (B) Quantification of polySer RAN aggregates in pons of non-transgenic and ATNX3^Q84/WT^ mice. ∗ p < 0.05; n=4/group for SCA3 YACQ84 and non-transgenic littermates. Red, positive staining; purple, nuclear counterstain.

Although these established mouse models have been used for decades to study the effects of polyGln, these mice also express both sense and antisense RAN proteins and the impact of these proteins, especially for the development of therapeutic strategies, needs to be considered.

### Locus-specific polySer and polyLeu RAN proteins are toxic to neural cells

The accumulation of polySer and polyLeu proteins in both grey and white-matter brain regions in human and mouse suggests RAN proteins are toxic to both neurons and glia. To test the toxicity of individual polySer and polyLeu RAN proteins in the absence of RAN translation or RNA gain of function effects caused by CAG_EXP_ or CUG_EXP_ RNAs, we generated alternative-codon minigenes with non-hairpin forming repeats that do not undergo RAN translation^11^. Each of these constructs contains an N-terminal Flag epitope tag, a polySer or polyLeu encoding repeat tract, and locus-specific (SCA1, SCA2, SCA3, SCA6, or SCA7) C-terminal regions unique to each of the polySer or polyLeu proteins (**Figure 7A**). These constructs were transfected into T98 (human glial), N2A (mouse neuroblast), and HEK293T (human embryonic kidney) cells and tested for cell toxicity by lactate dehydrogenase (LDH) assays 36 hours after transfection. Both polySer and polyLeu RAN proteins significantly increased cell death in T98 and N2A neural cells (**Figure 7B**). Additionally, polyLeu proteins were consistently more toxic to both N2A and T98 cells than the corresponding polySer protein for each disease (**Figure 7C**). A similar but not significant trend toward toxicity was seen with polySer in HEK293T cells. In contrast, toxicity was not seen for any of the SCA polyLeu proteins in non-neural HEK293T cells (**Figure 7B**).

**Figure 7.**
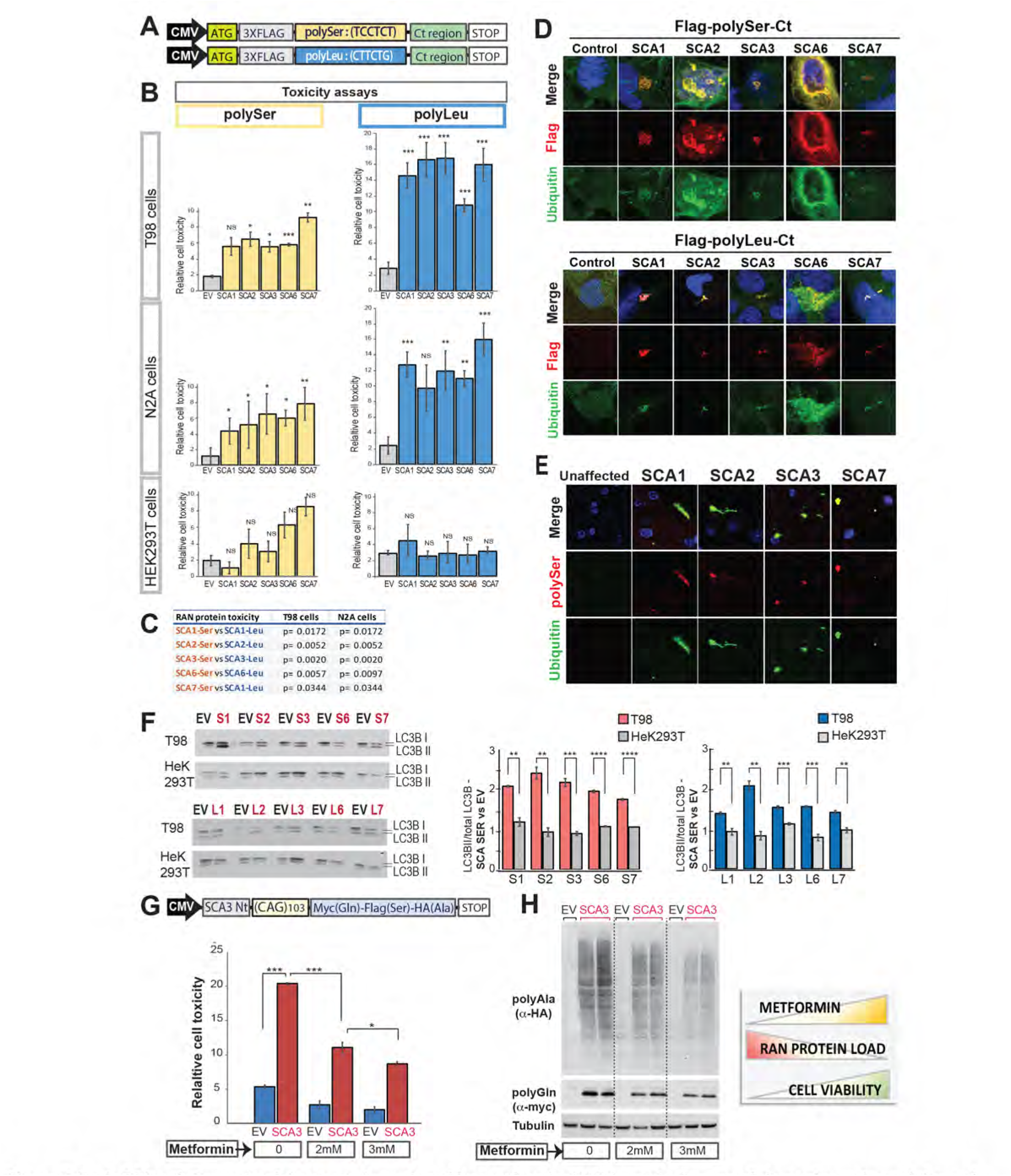
Toxic SCA polySer and polyLeu proteins reduced by metformin. (A) Schematic diagram of alternative codon polySer and polyLeu minigenes with predicted locus-specific C-terminal regions from SCA1, SCA2, SCA3, SCA6, and SCA7 expansion mutations. (B) Lactate dehydrogenase (LDH) assays showing percent cell death ± SEM (n=5) of T98, N2A and HEK293T cells expressing individual CAG-SCA polySer and polyLeu proteins. (C) Comparisons of locus specific polySer vs polyLeu toxicity in T98 and N2A cells (unpaired, two-tailed student’s t-test). (D) IF of transfected T98 cells showing colocalization of α-polySer and α-polyLeu (red) with α-ubiquitin (green) staining. (E) IF showing polySer RAN protein aggregates colocalize with ubiquitin staining in cerebellar white matter regions of SCA1, SCA2, SCA3, and SCA7. (F) Immunoblots show increased LC3B cleavage in T98 but not HEK293T cells expressing: SCA1-polySer (S1); SCA2-polySer (S2); SCA3-polySer (S3); SCA6-polySer (S6); SCA7-polySer (S7); SCA1-polyLeu (L1); SCA2-polyLeu (L2); SCA3-polyLeu (L3); SCA6-polyLeu (L6); SCA7-polyLeu (L7), with quantification in center and right panels, (G) Top Left: Non-ATG SCA3 CAG minigene used for transfections and LDH cytotoxicity assays and (H) RAN protein immunoblots of T98 cells expressing the SCA3 minigene +/- metformin treatment. n=4 independent experiments; ∗p < 0.05, ∗∗p < 0.01, ∗∗∗p < 0.001, ∗∗∗∗p < 0.0001.

In summary, these data demonstrate that SCA-derived polySer and polyLeu RAN proteins show cell-specific toxicity with increased cell death in T98 and N2A neural cells, which was not observed in HEK293T cells.

### SCA RAN protein aggregates colocalize with ubiquitin and alter autophagy

Immunofluorescence staining of T98 transfected cells shows that both polySer and polyLeu RAN proteins form defined cytoplasmic aggregates. SCA1, SCA2, SCA3, SCA6, and SCA7 polySer aggregates form round vesicle-like structures (**Figure 7D**). In addition, SCA6 polySer proteins can also accumulate as filamentous perinuclear aggregates (**Figure 7D**). For SCA1, SCA2, SCA3, and SCA7, the antisense polyLeu proteins form denser, more angular, and focal cytoplasmic aggregates. In contrast, SCA6 polyLeu proteins showed more abundant aggregates with both dense and diffuse staining (**Figure 7D**).

Ubiquitin-positive aggregates have been previously reported in white matter regions negative for polyGln in SCA1, SCA2, SCA3, and SCA7 but not in SCA6 autopsy brains. To test if RAN protein aggregates are ubiquitin-positive, we overexpressed SCA1, SCA2, SCA3, SCA6, and SCA7 polySer and polyLeu alternative-codon constructs in T98 cells. Immunofluorescence staining of these cells shows that SCA1, SCA2, SCA3, SCA6, and SCA7 polySer and polyLeu RAN protein aggregates colocalize with ubiquitin signal (**Figure 7D**). In human autopsy brains, IHC of RAN abundant and polyGln poor cerebellar white matter regions show polySer aggregates are ubiquitin-positive in SCA1, SCA2, SCA3, and SCA7 but not in SCA6 (**Figure 7E**). These results suggest that RAN proteins impair protein homeostasis.

Autophagic impairment has also been reported across different neurodegenerative disorders including the spinocerebellar ataxias^43–48^. To test the effects of SCA polySer and polyLeu RAN protein on autophagy, we compared LC3B cleavage in RAN vulnerable (T98) and more resistant (HEK293T) transfected cells. Immunoblotting shows polySer and polyLeu RAN proteins increase LC3BII levels relative to total LC3B in vulnerable T98 cells (**Figure 7F**), a change consistent with increased autophagosomes. In contrast, HEK293T cells, which are more resistant to polySer and polyLeu RAN proteins accumulation, showed no significant differences in LC3BII cleavage. Taken together, these results show that CAG-SCA RAN proteins alter key protein degradation pathways that are likely to contribute to the increased cell death seen in vulnerable neural cells.

### Metformin decreases SCA3 RAN protein levels and reduces toxicity in cultured cells

Metformin, an FDA-approved type-2 diabetes drug, has been shown to have beneficial therapeutic effects in both mice and humans in several repeat expansion disorders including FXS^49,50^, HD^51–54^, and DM1^55^. Recently, Zu et al.^21^ showed metformin reduces RAN protein levels and improves disease in C9*orf72* ALS/FTD BAC mice.

To test if metformin reduces RAN proteins and proteotoxicity in an SCA3 cell culture system, we transfected T98 cells with an SCA3 minigene. This construct contains 20 bp of endogenous human sequence upstream of the SCA3 repeat, an expanded CAG with 103 repeats, and three different Ct epitope tags to monitor RAN protein expression in all reading frames. (**Figure 7G**). Expression of a non-ATG SCA3 (CAG)_103_ minigene led to the accumulation of polyAla and polyGln RAN proteins and cell toxicity. Metformin-treated cells showed dose-dependent decreases in both RAN protein levels and cell toxicity (**Figure 7G-H**). These results suggest the potential utility of metformin as a repurposed drug to target RAN proteins expressed in CAG•CTG expansion disorders.

## DISCUSSION

Using both repeat and locus-specific C-terminal antibodies we show novel polySer and polyLeu RAN proteins, expressed from sense and antisense expansion transcripts respectively, form aggregates in SCA1, SCA2, SCA3, SCA6, and SCA7 autopsy brains. These proteins accumulate in regions of the brain most affected by these diseases including neurons and glia throughout the cerebellum and pons. While most studies of the polyGln SCAs have focused on degenerative changes in neurons, especially Purkinje cell neurons, recent reports have highlighted additional and extensive white matter abnormalities in these disorders^28–34^. White matter regions with frequent polySer and polyLeu RAN protein aggregates in the near absence of polyGln show striking pathological changes including demyelination, gliosis, and microglial activation. In contrast, polyGln aggregates are extremely rare in these white matter regions. Cell culture studies show SCA1, SCA2, SCA3, SCA6, and SCA7 polySer and polyLeu RAN proteins are toxic to both neuronal and glial cells. Additionally, polySer and polyLeu aggregates are ubiquitin positive and cause autophagy abnormalities. Inhibition of RAN translation using FDA-approved metformin decreases RAN protein levels and toxicity in an SCA3 cell culture model. Taken together, these data identify RAN translation as a common mechanism across the polyGln SCAs and suggest metformin may be a promising therapeutic strategy for the CAG-SCAs that warrants further research.

The polyGln SCAs are a prevalent group of ataxias that express elongated polyGln tracts as part of larger ATG-initiated open reading frames. Although there are currently no effective treatments to prevent or slow disease progression, multiple groups are focused on targeting the sense CAG^exp^ transcripts in these disorders^56–61^. While antisense oligonucleotide (ASO) strategies for SMA^62–64^ and SOD1^65–67^ ALS have been tremendously successful, unfortunately, ASO clinical trials for several repeat expansion diseases including HD^68^ and *C9orf72* ALS/FTD (https://investors.biogen.com/news-releases/news-release-details/biogen-and-ionis-announce-topline-phase-1-study-results) have failed. Although the ASO drugs used in each of these studies reduced proteins produced from targeted sense transcripts, both studies were halted because at the highest doses these drugs led to an increase of treatment related adverse events^68,69^. The biology of natural occurring transcripts is not fully understood but changes in levels of one RNA strand can affect the expression of the opposite strand through multiple mechanisms including RNA editing, RNA interference, nuclear localization, transcriptional interference, splicing alterations or translational regulation^70^. Previous studies in SCA7^3^ and HD^4^ have showed that sense and antisense transcripts regulate each other and that knockdown of one RNA results in the upregulation of the opposite strand transcript.

The overall strategy for the GAPmer ASOs used in the *C9orf72*^71^ and HD^68^ studies was to decrease levels of sense expansion transcripts by harnessing the power of the nuclear RNAse H pathway. ASO treatments that lower sense expansion transcripts may, however, disrupt the normal balance between sense and antisense expansion transcripts through one or more of the mechanisms described above. Moreover, changes to endogenous antisense transcripts may be difficult to detect because they are expressed at lower levels. Upregulation of naturally occurring antisense transcripts after ASO treatment in *C9orf72* ALS/FTD and HD could provide a molecular explanation for the worsening of the disease that was seen in patients treated with the highest doses of ASO^68,71^. Our observations that sense polySer and antisense polyLeu RAN proteins accumulate in CAG-SCA autopsy brains highlights the contributions of both sense and antisense mechanisms. Moreover, in each of the CAG-SCAs, antisense polyLeu RAN proteins are abundant in human brains and highly toxic to cells. Taken together, these data highlight the need to better understand the normal biology and contributions that naturally occurring antisense expansion transcripts play in repeat expansion diseases and to test strategies that reduce both sense and antisense expansion transcripts and/or RAN proteins.

White matter abnormalities have been reported across the polyGln SCAs^28–30^. MRI studies show white matter abnormalities are evident in early and premanifest stages of disease and are associated with motor and cognitive impairment in SCA2, SCA3 and SCA7^28–34^. The prominent accumulation of sense and antisense RAN proteins in near absence of polyGln in SCA1, SCA2, SCA3, SCA6 and SCA7 cerebellar white matter regions showing demyelination and neuroinflammatory changes, combined with the toxicity of polySer and polyLeu in T98 glial cells, suggests RAN proteins contribute to the white matter degeneration observed in these disorders. Similar white matter abnormalities are found across other CAG•CTG expansion disorders including HD, DM1 and SCA8. In HD and SCA8 RAN protein aggregates accumulate in similar white matter regions and are toxic to glial cells. These data link the molecular and cellular pathology of CAG•CTG expansion mutations across the polyGln SCAs with HD and the non-coding CAG•CTG disorders DM1 and SCA8.

An unresolved question in the polyGln field is what, at the molecular level, underlies selective patterns of neurodegeneration across diseases in which the CAG expansion RNAs and mutant polyGln proteins are broadly expressed across the body^2,72,73^. Cell culture experiments demonstrate that polySer and polyLeu proteins made from SCA1, SCA2, SCA3, SCA6, and SCA7 expansion mutations are toxic to neuronal and glial cells. In contrast, non-neural HEK293T cells are remarkably resistant to polySer and polyLeu overexpression. In the CAG-SCAs, RAN aggregates are primarily found in vulnerable regions of the cerebellum and brainstem with minimal accumulation in the relatively spared frontal cortex. In contrast, adult-onset HD cases show abundant RAN pathology in the more affected frontal cortex and less accumulation in the less involved cerebellum. The differences in RAN protein load, combined with the selective vulnerability of specific cells to RAN proteins, may contribute to the selective patterns of degeneration across these disorders.

Protein folding and degradation, which is important for cellular function, is compromised during aging^74^. The accumulation of misfolded proteins is a pathological hallmark of numerous age-related disorders, including Parkinson’s disease (PD), Alzheimer’s disease (AD) and HD^75^. Both autophagic impairment and ubiquitin inclusions in polyGln negative regions have been reported in the CAG-SCAs^43,45–47,76–78^. Our data showing that SCA polySer and polyLeu proteins colocalize with ubiquitin and alter autophagy in vulnerable cells suggests strategies that enhance autophagy and/or proteasomal activity may reduce RAN protein load and provide therapeutic benefit.

Metformin, a widely used FDA-approved drug for the treatment of type 2 diabetes can affect autophagy by interfering with the MAPK/mTOR signaling^79^, and its therapeutic potential in the treatment of neurodegenerative disorders has recently been explored^80^. A recent study shows that metformin decreases RAN protein accumulation across multiple types of repeat expansion mutations, likely by inhibiting the PKR pathway^21^. In C9 ALS/FTD BAC mice, metformin treatment reduced RAN protein levels, improved behavioral deficits, and increased motor neuron survival^21^. Metformin has also shown promise in HD and DM1^51–55^. These data, combined with our results showing decreased cell toxicity and SCA3 RAN protein levels, suggest metformin may be a promising therapeutic strategy for the CAG-SCAs that warrants further research.

The accumulation of toxic sense and antisense RAN proteins across the polyGln CAG-SCAs fundamentally shifts our understanding of these diseases. We show that each of these mutations, which were previously thought to encode a single mutant polyGln protein, express a cocktail of sense and antisense expansion proteins that accumulate in affected brain regions including damaged cerebellar grey and white matter regions. In addition to expanding the list of RAN protein diseases, the shared mechanism of RAN translation across numerous types of CAG•CTG diseases provides novel opportunities for drug and biomarker development across these disorders.

## Supporting information

Banez Coronel et al_Supplemental

## ACKNOWLEDGEMENTS

We thank the many ataxia patients and families for their generous contributions of autopsy tissue, the National Ataxia Foundation/University of Florida Center for NeuroGenetics Brain Bank, National Institutes of Health Grants R01NS098819, R37NS040389, R01NS117910; RF1NS098819, National Ataxia Foundation grant AGRDTD03-25-2022, Johns Hopkins Alzheimer’s Disease Research Center, NIH P30AG 066507; Dr. John Cleary for helpful discussions, Ms. Lisa A. Duvick and Dr. Harry T. Orr for generously providing mouse brain tissue from ATXN1 KI 145Q mice and their respective WT littermates, and Lauren Laboissonniere and Hannah Shorrock for technical assistance.

## METHODS

### Cell culture

T98 (human glioblastoma), HEK293T (human embryonic Kidney) and N2A (Neuro-2a, mouse neuroblast) cells were grown under standard conditions of temperature (37°C), humidity (95%), and carbon dioxide (5%) and cultured in Dulbecco’s Modified Eagle’s Medium (DMEM, Corning) supplemented with 10% FBS (Fetal Bovine Serum, Corning Cellgro), 100 units/ml penicillin and 100μg/ml Streptomycin (Corning Cellgro).

Metformin treatments were initiated 3 hours after transfection and were maintained for 33 hours until cell processing.

### DNA Constructs

*Flag-Ser-Ct and Flag-Leu-Ct* vectors described in^15^ were used for the optimization of polySer and polyLeu antibodies.

#### polySer, polyLeu alternative codon vectors

Minigenes for each polySer and polyLeu protein expressed from the SCA1, SCA2, SCA3, SCA6 and SCA7 repeat mutation were designed, synthesized by ADT Technologies, and inserted into p3XFLAG vectors. To avoid RNA hairpins and prevent RAN translation while encoding the same polySer or polyLeu homopolymeric motif, the CAG•CTG expansion was substituted by TCTTCC repeats in polySer-expressing constructs and CTTCTC repeats in polyLeu-expressing constructs. Each minigene contained a Flag tag at the Nt region, followed by the polySer or polyLeu repeat expansion and the unique Ct region for each protein. For SCA1, SCA2, SCA3 and SCA7 alternative codon constructs, minigenes contained seventy polySer or polyLeu repeats. SCA6 polySer and polyLeu constructs contained 30 repeats. The integrity of all constructs was confirmed by sequencing.

### Transfections

Transfection experiments were performed at a 60% cell confluence using Lipofectamine 2000 (Life technologies), according to the manufacturer’s protocol and with reduced DNA concentration (100ng of plasmid/cm^2^ of cultured cell monolayer). Cells were plated 24 hours before transfection. A GFP-expressing vector was used as a reporter to assess transfection efficiency 18 hours after transfection.

### Production of Polyclonal Antibodies

The polyclonal antibodies were generated by Biosynth. The α-polySer and α-polyLeu repeat antibodies were raised against synthetic peptides containing a stretch of (Ser)_10_ or (Leu)_12_ respectively.

Antisera for the Ct regions of antisera were generated with synthetic peptides from the C-terminal regions of the predicted polySer and polyLeu proteins translated from the SCA1, SCA2, SCA3, SCA6 and SCA7 repeat expansion mutations. The peptide sequence for each antibody epitope is: SCA1-Ser, RGSSPRGPPHQPSRT; SCA1-Le, SRRLAQTAHVGQQS; SCA2-Ser, RRPRLPMSASPAA; SCA2-Leu, GLQRHGEGPIHRLAR; SCA3-Ser, RGTYQDRVHIHVKGQ; SCA3-Leu, FFKVGFSSLPKLF; SCA6-Ser, TQAPRPSLWPEIGRPRGC; SCA6-Leu, RGPRAAGLTDHRGIGH; SCA7-Ser, RRLRSPSGSSTRHRRH; SCA7-Leu, LRHSFRSFSLEGRPK.

### Human Autopsy Tissue

SCA1, SCA2, SCA3, SCA6, SCA7, SCA5, HD and control autopsy tissues were collected at Johns Hopkins University, University of Minnesota and the University of Florida with informed consent of patients or their relatives and approval of respective institutional review boards.

### Mouse samples

Mice were housed in a specific pathogen-free facility, and animal care adhered to guidelines established by the Institutional Animal Care at the University of Florida, University of Michigan and University of Minnesota. SCA1 mouse tissue from the University of Minnesota was kindly provided by Lisa Duvick and Dr. Harry Orr.

### Immunohistochemistry

For the detection of SCA1, SCA2, SCA3, SCA6 and SCA7 polySer and polyLeu RAN proteins we used a triple-antigen retrieval protocol. Six-micrometer sections of paraffin-embedded tissue were deparaffinized in xylenes (2×15minutes) and rehydrated through an alcohol gradient (100%, 100%,95%, 80%; 10 minutes each). Sections were subsequently treated with the following antigen retrieval steps: A) 1ug/mL proteinase K treatment in 1mM CaCl2, 50mM Tris buffer (pH=7.6) for 45 minutes at 37°C. B) Pressure cooked in 10mM EDTA (pH=6.5) for ∼15minutes (until the pressure indicator lifts). C) 95% formic acid treatment for seven minutes^81,82^. Endogenous peroxidase block was performed in 3% H2O2 in methanol for twelve minutes followed by washings in running water for 20 minutes. To block non-specific binding, a non-serum block (Biocare Medical) was applied for 17 minutes at room temperature (RT).

Primary antisera were diluted in 1:10 non-serum block/water at the concentrations indicated below and incubated at 4^°^C overnight. Rabbit Linking Reagent (Covance) was applied for 35 minutes at RT. Secondary antibodies were Biotin-Avidin/Streptavidin labeled using ABC reagent (Vector laboratories, Inc.). Each step was followed by 3×5 minutes washes in PBS. RAN protein detection was performed by exposure to Vector Red Substrate Kit (Vector Laboratories, Inc.). Slides were washed in running water (10 mins) and counterstain was performed using Hematoxylin QS (Vector Laboratories) at RT (30 secs for human tissue, 2 mins for mouse tissue). After washing in water for 5 minutes, slides were dehydrated (ethanol 80%, 95%, 100%, xylene, 20 dips for each step) and mounted using Cytoseal 60 (Electron microscopy sciences). Images were captured with an Olympus BX51 light microscope.

Primary antibodies/sera were used at the following conditions in human postmortem tissue: α-polySer (1:5000, 5 min), α-polyLeu (1:6000, 6 min), α-Iba1 (Novus biological, NB100-1028, 1:1000, 3 min 30secs), 1C2 (Millipore, mouse, 1:10000, 1 min 30secs for striatum, 1:1000, 1min 30secs for cerebellum), α-SCA1-Ser-Ct (1:3000, 7min), α-SCA1-Leu-Ct (1:5000 8min), α-SCA2-Ser-Ct (1:6000 5min), α-SCA2-Leu-Ct (1:2500, 2min), α-SCA3-Ser-Ct (1:3000 7min), α-SCA3-Leu-Ct (1:7000, 6min), α-SCA6-Ser-Ct (1:3000 3min), α-SCA6-Leu-Ct (1:6000 3min30sec), α-SCA7-Ser (1:3000 5min), α-SCA7-Leu (1:7000 7min).

For mouse IHC, antibody concentrations were: α-polySer (1:8000, 4 min), α-polyLeu (1:8000, 6 min), α-SCA1-Ser-Ct (1:3000, 7min), α-SCA1-Leu-Ct (1:5000 8min), α-SCA3-Ser-Ct (1:10000 3min), α-SCA3-Leu-Ct (1:10000, 3min).

### Luxol fast blue staining

Paraffin-embedded sections (6μm) of human postmortem brain tissue were deparaffinized in xylene and hydrated to 95% ethyl alcohol, as conducted in immunohistochemistry experiments. The sections were then incubated in LFB solution (0.1% luxol fast blue in 95% ethyl alcohol, 0.5% acetic acid) at 60°C overnight. The next day, slides were rinsed in 95% ethyl alcohol (ten times) and distilled water (3mins). Subsequently, the slides were differentiated in 0.05% lithium carbonate solution for 5secs and 70% ethyl alcohol (2x 10 sec) and washed in distilled water. The slides were dehydrated through ethanol 80%, 95%, 100%, 100%, 20 dips for each step; and xylene (3mins x 2); and mounted using Cytoseal 60 (Electron microscopy sciences) before visualization. Images were captured with an Olympus BX51 light microscope.

### Immunofluorescence

T98 and HEK293T cells were grown on coverslips and transfected as previously described. Cells were processed 42 hours after transfection for antibody optimization experiments, and 24 hours after transfection for Ubiquitin colocalization experiments. Cells were washed with PBS, fixed with 4% paraformaldehyde in PBS (20 min at RT), rinsed in PBS and permeabilized with 0.5% Triton-X-100 in PBS for 30 min at RT. Non-specific binding was blocked by incubation in 10% normal goat serum (NGS) in PBS for 1 hour at RT. The indicated primary antibodies were diluted in 1% NGS in PBS, and incubated overnight at 4°C. After washing (3x 5min) in PBS 1X, coverslips were incubated with secondary anti-mouse IgG Alexa 488, anti-rabbit IgG Alexa 594 (ThermoFisher) or anti-sheep IgG (Vectorlabs) at a dilution of 1:2000 for 1 hour at RT.

Coverslips were washed, incubated with DAPI (0.1 μg/ml in PBS, 15 min), washed once and mounted in Faramount Aqueous mounting medium (Dako). Cells were visualized under a Zeiss confocal microscope. Primary antibodies were anti-Flag M2 (1:1500, Sigma), α-Ser-Ct (1:500), α-Leu Ct (1:1500), α-polySer (1:500), α-polyLeu Ct (1:1000), α-Ubiquitin (1:300, Sigma).

Immunofluorescence on human postmortem fixed tissue was conducted following day 1 protocol of immunohistochemistry assays and day 2 of immunofluorescence in fixed cells. Primary antibodies were used at 1:4000, for α-polySer, and 1:500 for α-Ubiquitin. Sudan black was applied to block autofluorescence.

#### Immunoblotting

Cells in six-well tissue-culture plates were rinsed with PBS and lysed in 80 μL RIPA buffer for 5 minutes on ice. The cell lysates were collected and centrifuged at 14,000 x g for 17min at 4°C. The protein concentration of the supernatant was determined using the protein assay dye reagent (Bio-Rad). Thirty micrograms of protein were separated in a 4-12% NuPage Bis-Tris gel (Invitrogen) and transferred to a nitrocellulose membrane (Amersham). The membrane was blocked in PBS containing 0.05% Tween-20 detergent (PBS-T) and 5% non-fat dry milk powder. Primary antibodies were prepared in 1% milk in PBS-T and incubated overnight at 4°C. After washing with PBS-T (3×15min), membranes were incubated with secondary antibodies for 1 hour at RT and washed with PBS-T (3×10min). Detection was performed using Western Lightning Plus-ECL (Perkin Elmer). Primary antibodies were α-LC3B (PA1-16930, 1:2000, Thermofisher) and α-GAPDH (1:5000, ab8245 Abcam). Peroxidase-conjugated anti-rabbit secondary antibodies were used at 1:2000 (GE Healthcare).

### Cell-Toxicity Assays

Cell toxicity was determined measuring lactate dehydrogenase release from dying cells using Cytotox 96 non-radiactive assay (Promega), following manufacturer’s instructions. LDH determinations were performed in five independent experiments for alternative codon studies and four independent experiments for metformin toxicity studies, with each experiment performed in quintuplicates. Measurements were conducted at 490nm.

### Statistical analysis

Statistical significance was calculated using the two-tailed unpaired t-student’s test for single comparisons (p<0.05). The analysis of variance (ANOVA) was used for the comparison of multiple pairwise conditions. “n” refers to independent experiments.

