## Supplementary material for "Repeat associated non-AUG translation as a common mechanism for the polyGln ataxias": Banez Coronel et al_Supplemental

[1]Center for NeuroGenetics; [2]Dept. of Molecular Genetics and Microbiology, College of Medicine, University of Florida, Gainesville, FL; [3]Dept. of Pathology and Anatomical Sciences, University at Buffalo the State University of New York, NY; [4]Dept. of Pathology, Immunology and Laboratory Medicine, College of Medicine, University of Florida; [5]Dept. of Pathology, John Hopkins University School of Medicine, Baltimore, MD; [6]Dept. of Neurology, University of Michigan, MI; [7]Dept. of Neurology, Houston Methodist Hospital, TX; [8]McKnight Brain Institute, University of Florida; [9]Department of Neurology, College of Medicine, University of Florida; [10]Norman Fixel Institute for Neurological Diseases, University of Florida, Gainesville, FL.[11]Genetics Institute, University of Florida; [12]Lead Contact

*Corresponding authors

Lead Contact:

Laura P.W. Ranum, Ph.D.

Director, Center for NeuroGenetics

Professor of Molecular Genetics and Microbiology, College of Medicine

University of Florida

2033 Mowry Road

PO Box 103610

Gainesville, FL 32610-3610

352 294-5209

| Table S1. Summary of SCA and control cases | | | | |
|---|---|---|---|---|
| | AOD | Sex | Race | PMI (hours) |
| SCA1 A | 42 | M | N/A | 7 |
| SCA1 B | 80 | M | W | 12 |
| SCA1 C | 46 | M | N/A | 55 |
| SCA1 D | 64 | M | W | 9.5 |
| SCA1 E | NA | N/A | N/A | N/A |
| SCA2 A | 66 | F | N/A | 3.5 |
| SCA2 B | 60 | F | N/A | 16.5 |
| SCA2 C | N/A | N/A | N/A | N/A |
| SCA2 D | 74 | F | W | 19 |
| SCA2 E | 51 | M | N/A | 25 |
| SCA3 A | 44 | M | N/A | 15 |
| SCA3 B | 55 | M | N/A | 5 |
| SCA3 C | 66 | F | N/A | 18 |
| SCA3 D | 49 | M | N/A | 26 |
| SCA3 E | 62 | F | W | 37 |
| SCA3 F | 48 | F | B | 8.5 |
| SCA6 A | 74 | F | N/A | 19 |
| SCA6 B | 85 | F | NR | 7.75 |
| SCA6 C | N/A | N/A | N/A | N/A |
| SCA6 D | 88 | F | N/A | 24 |
| SCA7 A | 69 | F | N/A | 21.5 |
| SCA7 B | 27 | M | N/A | N/A |
| SCA7 C | 48 | F | N/A | 24-48 |
| SCA7 D | 28 | M | N/A | N/A |
| SCA7 E | 78 | F | N/A | 48 |
| SCA5 A | NR | M | N/A | N/A |
| SCA5 B | N/A | N/A | N/A | N/A |

**Table S1. Summary of SCA and control cases.** AOD, age of death; N/A, not available; M, male; F, female; PMI, post-mortem interval.

**Table S2 Summary of RAN protein staining using polySer and polyLeu Ct antibodies in the polyGln SCAs**

| | α-polySer | α-SCA1-Ser-Ct | α-SCA2-Ser-Ct | α-SCA3-Ser-Ct | α-SCA6-Ser-Ct | α-SCA7-Ser-Ct |
|---|---|---|---|---|---|---|
| **SCA1** (n=5) | +++ | +++ | - | - | - | - |
| **SCA2** (n=5) | ++++ | - | +++ | - | - | - |
| **SCA3** (n=6) | +++ | - | - | ++++ | - | - |
| **SCA6** (n=4) | ++++ | - | - | - | ++++ | - |
| **SCA7** (n=5) | ++++ | - | - | - | - | +++ |
| **CRTL** (n=5) | - | - | - | - | - | - |
| | α-polyLeu | α-SCA1-Leu-Ct | α-SCA2-Leu-Ct | α-SCA3-Leu-Ct | α-SCA6-Leu-Ct | α-SCA7-Leu-Ct |
| **SCA1** (n=5) | +++ | +++ | +/- | - | - | - |
| **SCA2** (n=5) | ++++ | - | ++++ | - | - | - |
| **SCA3** (n=6) | +++ | - | +/- | +++ | - | - |
| **SCA6** (n=4) | ++++ | - | +/- | - | ++++ | - |
| **SCA7** (n=5) | ++++ | - | +/- | - | - | ++++ |
| **CTRL** (n=5) | - | - | - | - | - | - |

**RAN protein staining:** (-) no signal; (+/-) rare α-SCA2-Leu-Ct signal seen in a subset of genetically confirmed ataxia cases negative for SCA2 [1/5 SCA1, 1/6 SCA3, 1/4 SCA6 and 1/5 SCA7 cases]; (+++) frequent intense staining in all cases, ++++ very frequent and highly intense staining in all cases.

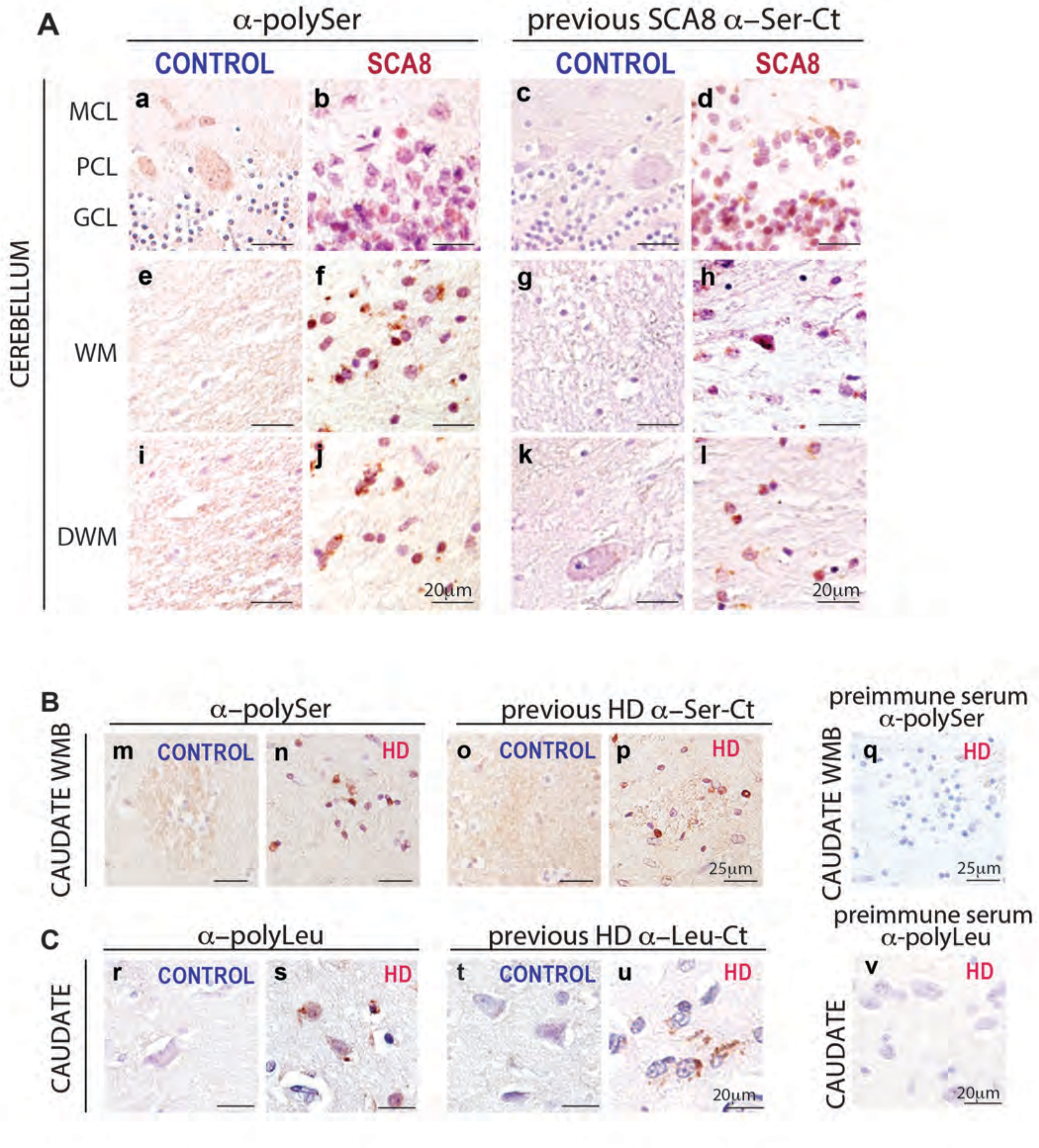


**Supplementary Figure 1. Novel polySer and polyLeu repeat antibodies specifically detect RAN protein aggregates in HD and SCA8 brain.** (A) IHC showing newly generated ⊠-polySer repeat motif antibody detects aggregates in the granular cell layer (a,b), cortical white matter (e,f) and deep white matter regions (I,j) of SCA8 but not control cerebellum, similar to SCA8 polySer aggregates detected with the previously validated SCA8-Ser-Ct antibody (d,h,l). Control tissue did not show similar signal (c,g,k). (B, C) Large field images of IHC staining show similar nuclear signal in HD striatal regions using ⊠-polySer (n) and ⊠-polyLeu (s) antibodies compared to previously validated HD-Ser-Ct and HD-Leu-Ct antibodies[13] (p,u), respectively. No similar signal was detected in control samples or HD tissue stained with pre-immune sera (q,v). Red, positive staining; blue, nuclear counterstain; GCL, granular cell layer; PCL, Purkinje cell layer; MCL, molecular cell layer; WM, white matter; DWM, deep white matter; WMB, white matter bundles.

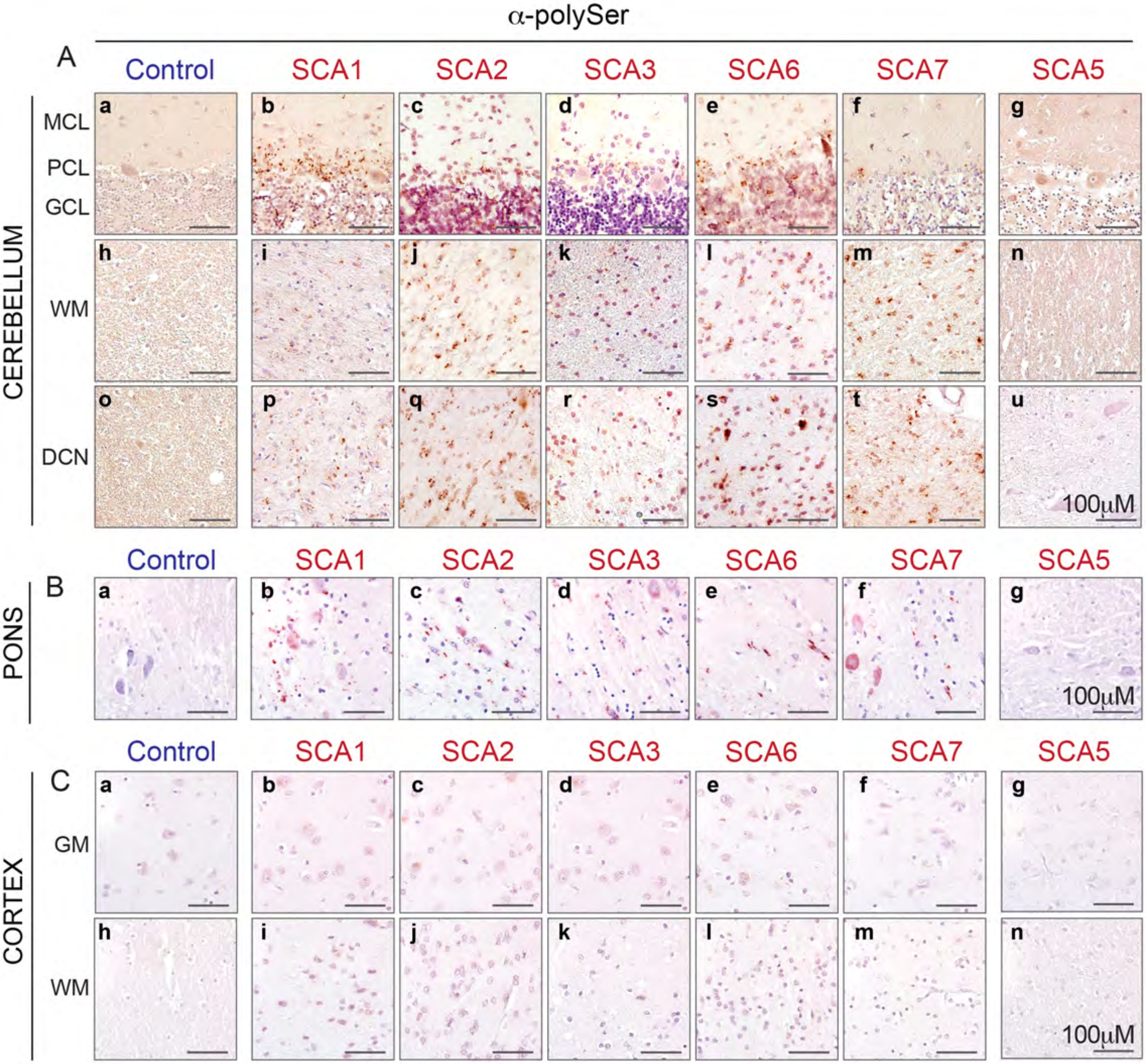


**Supplementary Figure 2. RAN-polySer proteins accumulate as frequent and intense aggregates in SCA1, SCA2, SCA3, SCA6, and SCA7 cerebellum and pons.** Large field images of ⊠-polySer IHC staining show prominent accumulation of nuclear, perinuclear and neuropil aggregates throughout the cerebellum in SCA1, SCA2, SCA3, SCA6 and SCA7 (b-f, i-m, p-t) but not unaffected (a,h,o) or disease controls (g,n,u). (B) Frequent ⊠-polySer aggregates in pons across the CAG-SCAs in neurons and glia (b-f) but not controls (a, g). (C) Minimal ⊠-polySer signal is found across the CAG-SCAs in the frontal cortex (b-f, i-m). GCL, granular cell layer; PCL, Purkinje cell layer; MCL, molecular cell layer; WM, white matter; DCN, deep cerebellar nuclei; GM grey matter. Red, positive ⊠-polySer staining; purple, nuclear counterstain.

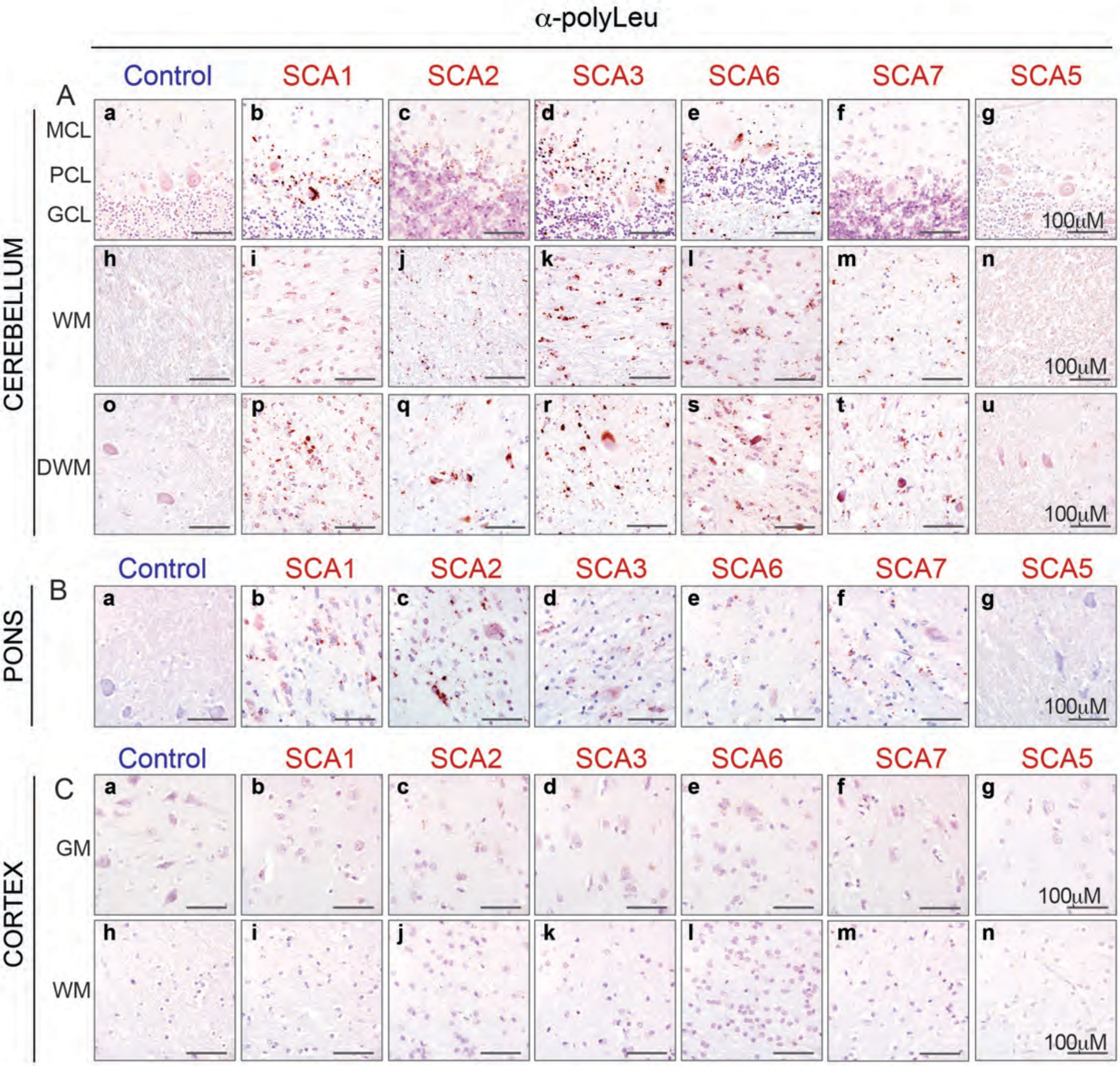


**Supplementary Figure 3. Antisense RAN protein aggregates throughout SCA1, SCA2, SCA3, SCA6, and SCA7 cerebellum and pons.** Large field images of $\alpha$-polyLeu IHC staining show prominent accumulation of nuclear, perinuclear and neuropil aggregates throughout the cerebellum in SCA1, SCA2, SCA3, SCA6 and SCA7 (b-f, i-m, p-t) but not unaffected (a,h,o) or disease controls (g,n,u). (B) Frequent $\alpha$-polyLeu aggregates seen in neurons and glia throughout the pons across the CAG-SCAs (b-f). (C) Minimal $\alpha$-polyLeu signal is found across the CAG-SCAs in the frontal cortex (b-f, i-m). Red, positive $\alpha$-polyLeu staining; purple, counterstain; GCL, granular cell layer; PCL, Purkinje cell layer; MCL, molecular cell layer; WM, white matter; DWM, deep white matter; GM grey matter.

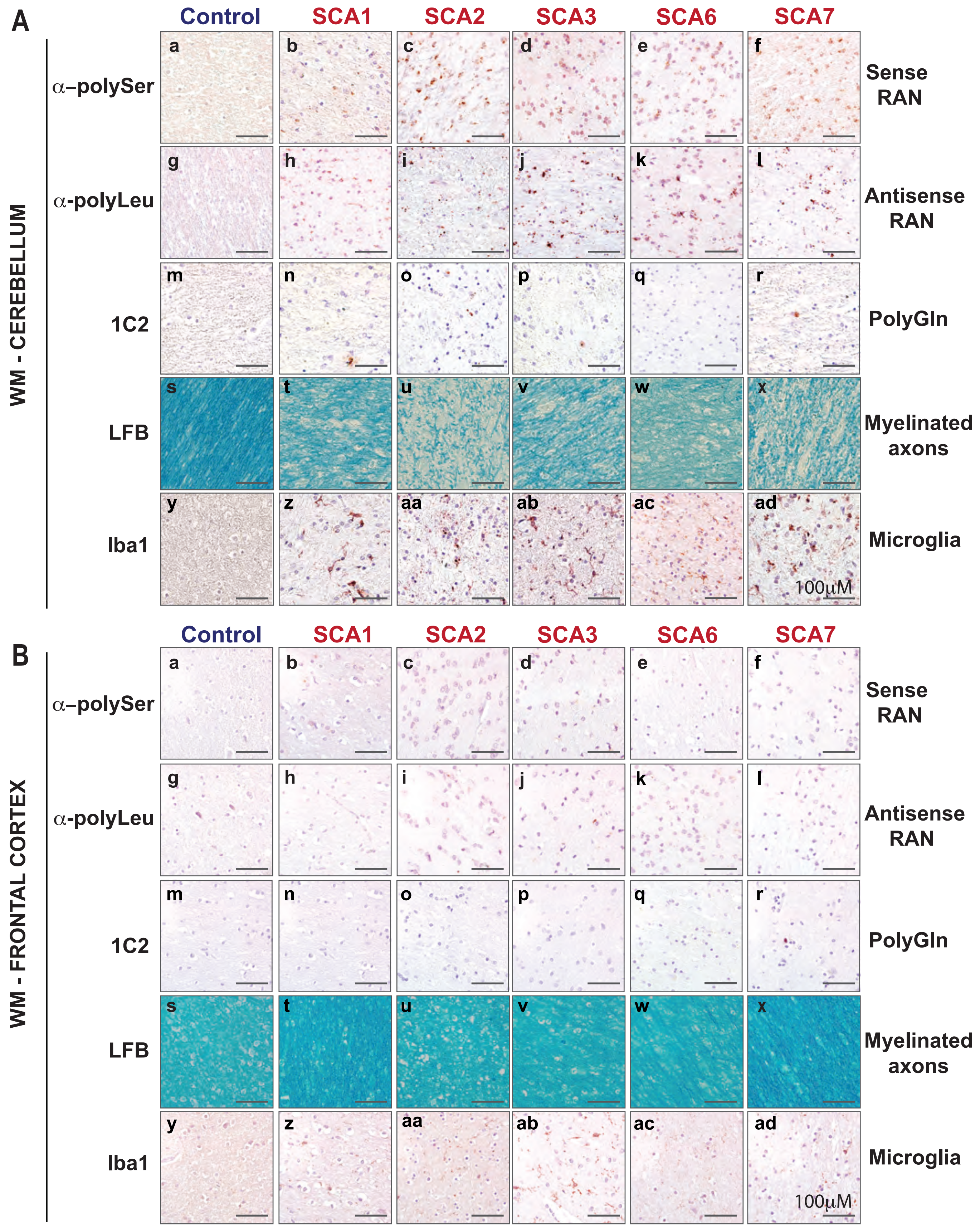


**Supplementary Figure 4. Abundant RAN protein aggregates in cerebellar white matter regions with pathological changes and only rare polyGln.** (A) Large field IHC images of SCA1, SCA2, SCA3, SCA6 and SCA7 cerebellum show frequent RAN polySer (a-f) and RAN polyLeu aggregates (g-l) and rare polyGln aggregates (m-r) in white matter regions with prominent demyelination, shown by decreased luxol fast blue (LFB) signal (s-x) and increased microglia (y-ad) compared to controls. (B) In contrast, white matter regions in the frontal cortex show minimal RAN protein staining, no increase in Iba1 signal and no obvious alterations in LFB intensity compared to controls. Luxol Fast Blue (LFB), WM, white matter. Red, positive staining; purple, nuclear counterstain. Control, n=5; SCA1, n=5; SCA2, n=5; SCA3, n=6; SCA6, n=4; SCA7, n=5.

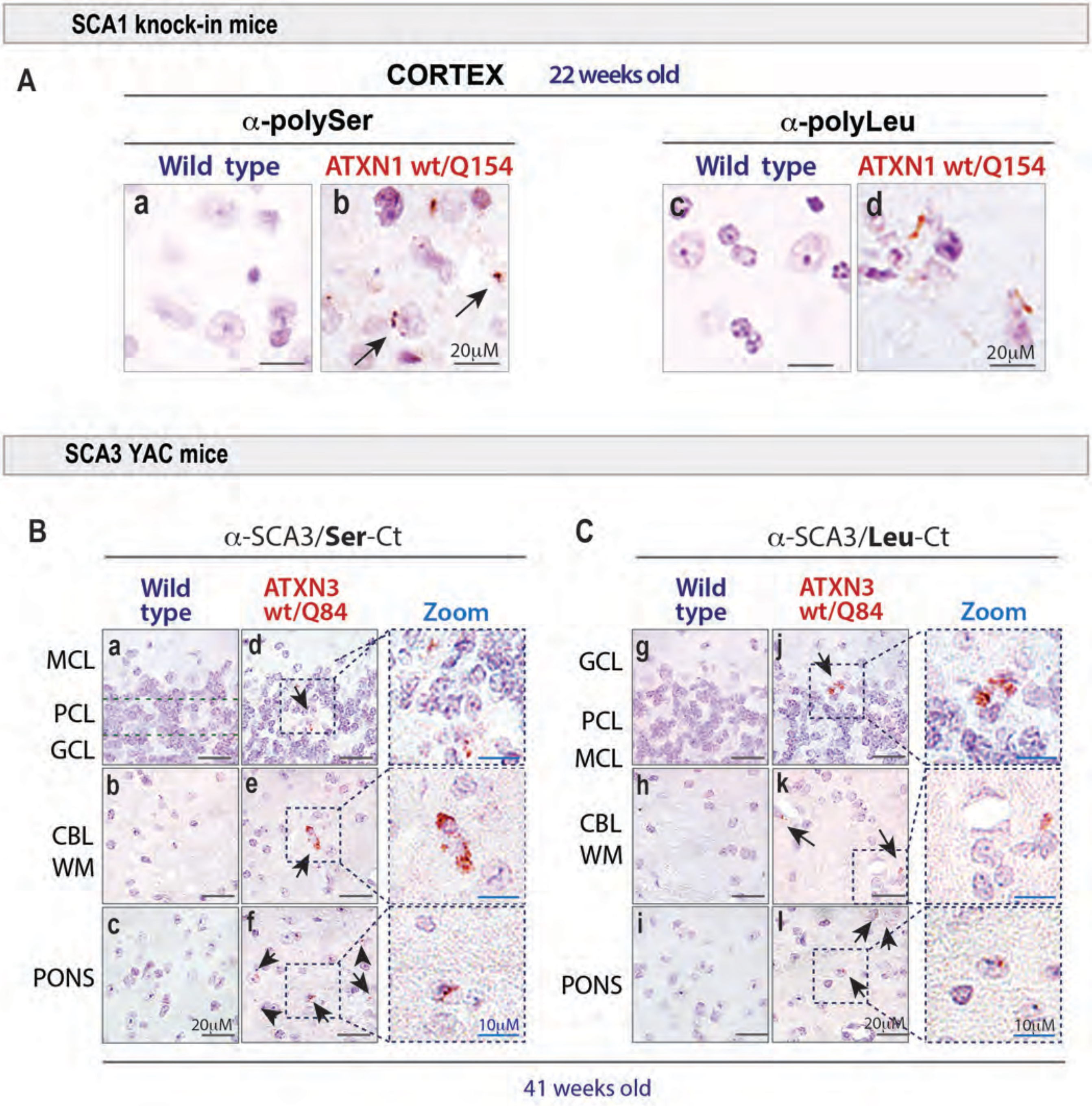


**Supplementary Figure 5. RAN proteins accumulate in SCA1 and SCA3 ataxia mouse models.** (A) IHC staining using novel ⊠-polySer and ⊠-polyLeu repeat antibodies detect sense and antisense RAN protein aggregates in the cortex of SCA1 ATXN1154Q/WT knock-in mice. (B-C) IHC with ⊠-SCA3-Ser-Ct and ⊠-SCA3-Leu-Ct antibodies detect aggregates in the granular cell layer, cerebellar white matter regions and pons from SCA3 YAC ATNX3Q84/WT. n=3/group for SCA1 Q154 mice and non-transgenic littermates; n=4/group for SCA3 YACQ84 and non-transgenic littermates. GCL, granular cell layer; WM, white matter; Red, positive staining; purple, nuclear counterstain.
